# Neurotensin neurons in the central extended amygdala control energy balance

**DOI:** 10.1101/2021.08.03.454970

**Authors:** Alessandro Furlan, Alberto Corona, Sara Boyle, Radhashree Sharma, Rachel Rubino, Jill Habel, Eva Carlotta Gablenz, Jacqueline Giovanniello, Semir Beyaz, Tobias Janowitz, Stephen D. Shea, Bo Li

**Affiliations:** Cold Spring Harbor Laboratory, Cold Spring Harbor, NY 11724, USA; School of Biological Sciences, Cold Spring Harbor Laboratory, Cold Spring Harbor, NY 11724, USA; Medical Faculty, Ruprecht-Karls-University Heidelberg, Heidelberg, Germany

## Abstract

Overeating and a sedentary life style are major causes of obesity and related metabolic disorders. Identification of the neurobiological processes that regulate energy balance will facilitate development of interventions for these disorders. Here we show that the Neurotensin-expressing neurons in the mouse IPAC (IPAC^Nts^), a nucleus of the central extended amygdala, bidirectionally coordinate hedonic feeding and physical activity, thereby regulating energy balance, metabolic processes and bodyweight. IPAC^Nts^ are preferentially activated by consumption of highly palatable food or exposure to its taste and smell. Activating IPAC^Nts^ promotes food intake in a palatability-dependent manner and decreases locomotion. Conversely, inhibiting IPAC^Nts^ selectively reduces palatable food intake and dramatically enhances physical activity and energy expenditure, and in parallel stimulates physiological responses that oppose diet-induced obesity and metabolic dysfunctions. Thus, a single neuronal population, Neurotensin-expressing neurons in the IPAC, acts to control obesogenic and leptogenic processes by synergistically coordinating energy intake and expenditure with metabolism.

## INTRODUCTION

In past decades obesity has become an epidemic and is currently one of the main causes of premature death worldwide (Global et al., 2016; Mitchell et al., 2011). Genetic, environmental, and behavioral factors all contribute to the onset and progression of weight gain and obesity. Overeating and sedentary behavior, which are common in modern societies, support a positive energy balance and in the long-term lead to weight gain and metabolic disorders. Treatments involving lifestyle changes with the goal of losing weight and ameliorating metabolic diseases often fail, in part because metabolic adaptations following weight loss act to restore homeostasis and the original body weight (Fothergill et al., 2016; Trexler et al., 2014).

Homeostasis is regulated by specialized homeostatic neurons in the brain located in the hypothalamus, parabrachial nucleus (PBN), and the nucleus tractus solitarii (NTS), which receive orexigenic and anorexigenic inputs from the periphery. These neurons regulate energy intake via homeostatic feeding (i.e., feeding on the basis of metabolic need), and regulate energy expenditure to meet metabolic demands and maintain a stable body weight (Roh et al., 2016; Rossi and Stuber, 2018; Saper et al., 2002; Sternson and Eiselt, 2017).

However, palatable foods can elicit feeding in the absence of a metabolic need, a phenomenon known as hedonic eating (Morales and Berridge, 2020; Rossi and Stuber, 2018). In humans, the degree of food intake positively correlates with food palatability – the hedonic evaluation of food sensory cues such as smell, taste, and texture (Yeomans, 1998; Yeomans and Wright, 1991). Foods rich in carbohydrates and fat are usually highly palatable (DiFeliceantonio et al., 2018). As industrialization leads to an abundance of such highly palatable and calorie-dense foods, hedonic eating, which results in excessive energy intake, is considered a major contributor to the obesity epidemic.

In addition to excessive energy intake, insufficient energy expenditure is another major contributor to the development of obesity (Hill et al., 2012; Tremblay and Willms, 2003), as unused energy is stored in the body in the form of white adipose tissue (WAT) (Aldiss et al., 2018). The main channels of energy expenditure are adaptive thermogenesis, which is regulated by brown adipose tissue (BAT) activation in response to a thermal stimulus (e.g., cold temperature), and activity-dependent thermogenesis (Srivastava and Veech, 2019). Strenuous physical activity can induce the appearance of interspersed brown-like (beige) adipocytes in WAT. These cells, unlike white adipocytes, are metabolically active and dissipate energy (Aldiss et al., 2018; Srivastava and Veech, 2019).

While the neural circuits underlying homeostatic energy intake have been extensively characterized (Sternson and Eiselt, 2017), those regulating hedonic feeding and energy expenditure are less well understood (Gong et al., 2020; Hardaway et al., 2019; Riera et al., 2017; Schneeberger et al., 2019; Zhang and van den Pol, 2017). Substantial evidence indicates that structures belonging to the central extended amygdala (EAc), in particular the central amygdala (CeA) and the bed nucleus of the stria terminalis (BNST), play important roles in the maintenance of homeostasis (Cai et al., 2014; Douglass et al., 2017; Hardaway et al., 2019; Jennings et al., 2013; Wang et al., 2019). The interstitial nucleus of the posterior limb of the anterior commissure (IPAC) is another major structure of the EAc (Alheid, 2003). However, unlike the BNST and CeA, which have been intensively studied in the context of motivated behaviors including feeding, the function of the IPAC in behavior is largely unknown.

The IPAC encompasses a corridor of neurons extending from the caudal portion of the nucleus accumbens (NAc) shell, merging into the lateral nuclei of the BNST (rostral IPAC) and further reaching towards the CeA (caudal IPAC) (Alheid, 2003). Its afferent and efferent connections are grossly similar to those of the lateral BNST and the CeA (Alheid et al., 1999; Gehrlach et al., 2020; Shammah-Lagnado et al., 2001). For example, neurons in the IPAC receive dense projections from the gustatory insular cortex and in turn project to the lateral hypothalamus (LH), paraventricular nucleus of the hypothalamus (PVN), ventral tegmental area (VTA), NTS, and other brainstem areas, which are structures implicated in regulating energy balance, autonomic responses and reward processing (Sternson and Eiselt, 2017). Thus, the IPAC is anatomically poised to participate in regulating energy intake and/or expenditure.

It was recently shown that IPAC neurons are activated by innate or learned gustatory stimuli (Tanaka et al., 2019, 2021). Nevertheless, the IPAC has a unique narrow elongated shape and unclear anatomical borders with the rostral NAc and caudal BNST, making it difficult for targeted *in vivo* manipulation based only on its anatomy. One approach to address this issue is to use genetic markers to selectively label and study specific populations of IPAC neurons. Previous studies indicate that the neuropeptide Neurotensin (Nts) is expressed and enriched in the rostral IPAC region (Schroeder et al., 2019; Woodworth et al., 2018). Interestingly, Nts infusion into specific brain areas, including the PVN, VTA, and NTS, modulates feeding behavior, body temperature, and physical activities (for a recent review, see (Ramirez-Virella and Leinninger, 2021)). These findings indicate that central Nts has a role in regulating both energy intake and energy expenditure, and further suggest that Nts-expressing IPAC neurons may contribute to similar or related processes, as many of the Nts infusion areas are also the projection targets of IPAC neurons.

In this study, we characterized the function of Nts-expressing neurons in the rostral IPAC, and uncovered a critical role of these neurons in bidirectional control of energy intake and expenditure, thereby regulating metabolism and body weight.

## RESULTS

### IPAC^Nts^ neurons are specifically activated by palatable food

To verify Nts expression in the IPAC, we bred *Nts^Cre^;Ai14* mice, in which Nts-expressing (Nts^+^) neurons express the red fluorescence protein tdTomato (Leinninger et al., 2011; Madisen et al., 2010). We found that dense Nts^+^ cells form a narrow stripe in the most medial portion of the rostral IPAC (Figure 1A, Figure S1A), which merge with the sparser Nts^+^ cells in the lateral ventral (STLV) and juxtacapsular (STLJ) nuclei of the lateral BNST, forming a continuum. No Nts^+^ neurons were observed in the ventral pallidum (VP), though it is rich in axon fibers originating from Nts^+^ neurons (Figure 1A, Figure S1A).

**Figure 1.**
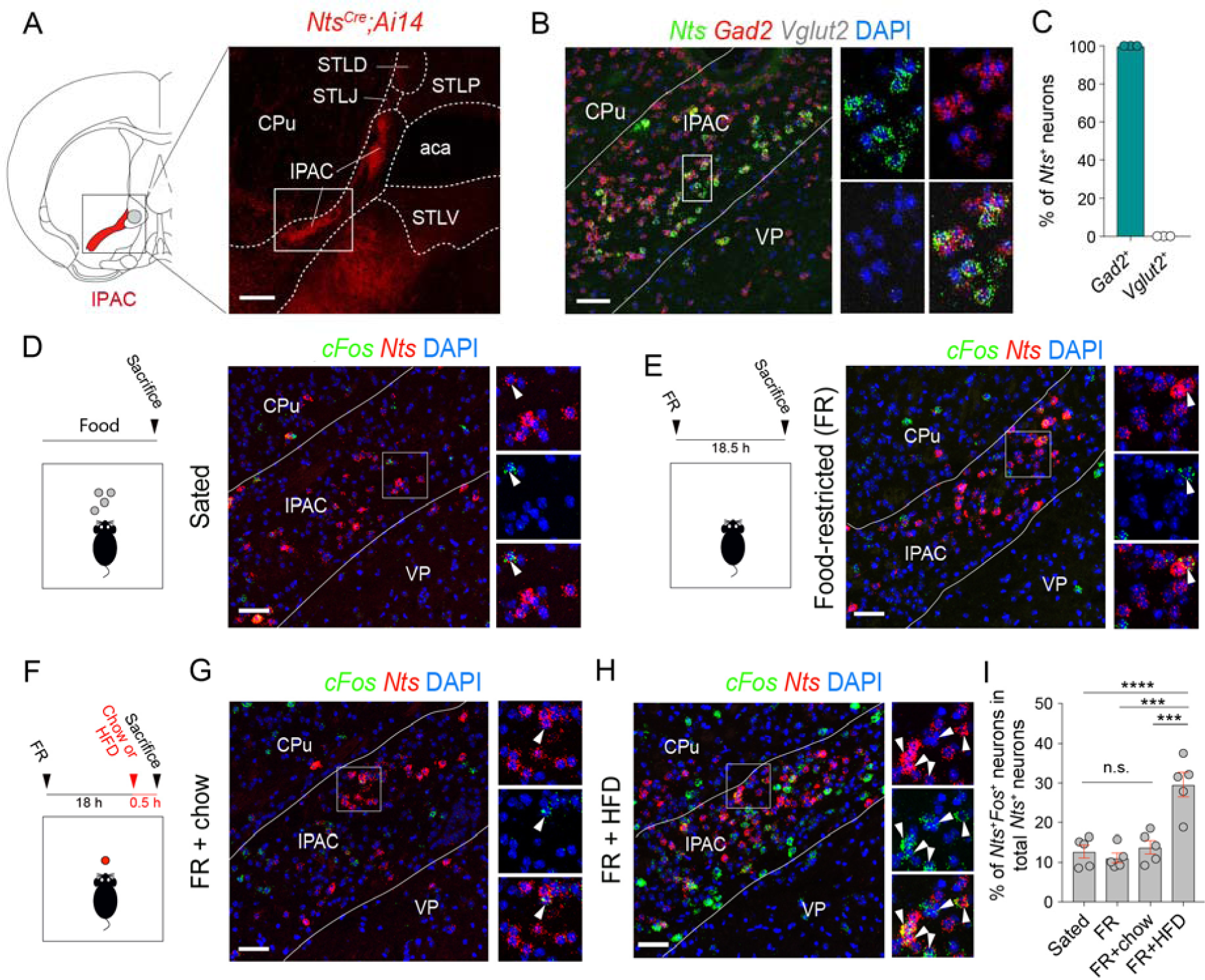
IPAC^Nts^ neurons are selectively activated by palatable food. (A) Left: a schematic of a coronal brain section containing the IPAC (red). Right: a confocal image of a coronal brain section from a representative *Nts^Cre^;Ai14* mouse, showing the distribution of Nts neurons in the IPAC (red). Scale bar: 200 µm. aca, anterior commissure; STLP/STLV/STLD/STLJ, lateral posterior/ventral/dorsal/juxtacapsular division of the bed nucleus of the stria terminalis; CPu, caudoputamen; VP, ventral pallidum. (B) Left: a representative confocal image of *in situ* hybridization for *Nts*, *Gad2* and *Vglut2*, and DAPI staining for nuclei. Right, high-magnification images of the boxed area on the left, showing *Nts*^+^ cells in the IPAC expressed *Gad2* but not *Vglut2*. Scale bar: 50 μm. (C) Quantification of the result in (B) (n = 3 mice). (D) Left: a schematic of the approach. Right: representative confocal images of *in situ* hybridization for *cFos* and *Nts* in the brain sections from sated mice. On the rightmost are high-magnification images of the boxed area, showing only few *Nts*^+^ cells in the IPAC expressed *cFos* (arrow heads). Scale bar: 50 μm. (E) Same as (D), except that the result was from food-restricted (FR) mice. (F) A schematic of the approach. (G) Representative confocal images of *in situ* hybridization for *cFos* and *Nts* in the brain sections from FR mice just fed with chow. On the right are high-magnification images of the boxed area, showing only few *Nts*^+^ cells in the IPAC expressed *cFos* (arrow heads). Scale bar: 50 μm. (H) Same as (G), except that the result was from FR mice just fed with HFD, and that many *Nts*^+^ cells in the IPAC expressed *cFos*. (I) Quantification of the results in (D-H). N = 5 mice in each group; F_(3,16)_ = 17.51, p < 0.0001; ***p < 0.001, ****p < 0.0001; one-way ANOVA followed by Sidak’s multiple comparisons test. Data are presented as mean ± s.e.m.

Single molecule fluorescent *in situ* hybridization (smFISH) confirmed *Nts* expression in the IPAC and lateral nuclei of the BNST, and its near absence in nearby striatal and VP territories. Virtually all *Nts*^+^ neurons in the IPAC (hereafter referred to as IPAC^Nts^ neurons) were GABAergic (Figure 1B, C; Figure S1B). In addition, the expression pattern of *Cre* recapitulated that of endogenous *Nts* in the IPAC of *Nts^Cre^* mice (Figure S1C), thus validating the fidelity of this line.

Food-restriction creates a negative energy balance and leads to the activation of homeostatic circuits regulating energy intake to restore the balance (Atasoy et al., 2012). To test whether IPAC^Nts^ neurons are involved in this process, we analyzed the expression of *c-Fos*, a marker for neuronal activation, in these neurons in food-restricted (FR) or sated mice. We found that food-restriction did not alter the *c-Fos* expression (Figure 1D, E, I).

Next, we exposed food-restricted mice to either regular chow or a high-fat diet (HFD) (Methods). Interestingly, feeding on the HFD, but not regular chow, induced a marked increase in *c-Fos* expression in IPAC^Nts^ neurons (Figure 1F-I; Figure S1D, E), although the mice consumed comparable amounts of chow and HFD (Figure S1F, left) and had similar energy intake (Figure S1F, right). In contrast, feeding on either the regular chow or HFD induced robust *c-Fos* expression in the adjacent striatum (Figure S1D, E). These results suggest that IPAC^Nts^ neurons are activated preferentially by the consumption of palatable food, but not by an energy deficit or food consumption *per se*.

### IPAC^Nts^ neurons encode tastant palatability

We reasoned that IPAC^Nts^ neurons encode food palatability. To test this hypothesis, we set out to monitor the *in vivo* activities of these neurons in behaving mice consuming substances with differing palatability. We first labelled these neurons with the genetically encoded calcium indicator GCaMP6 (Chen et al., 2013), by injecting the IPAC of *Nts^Cre^* mice with an adeno-associated virus (AAV) expressing GCaMP6f in a *Cre*-dependent manner, then implanting an optical fiber into the same location (Figure 2A, B, Figure S2A). This strategy allows recording bulk GCaMP6 signals, which are readouts of average neuronal activities, *in vivo* from the infected neurons with fiber photometry (Xiao et al., 2020; Yu et al., 2016).

**Figure 2.**
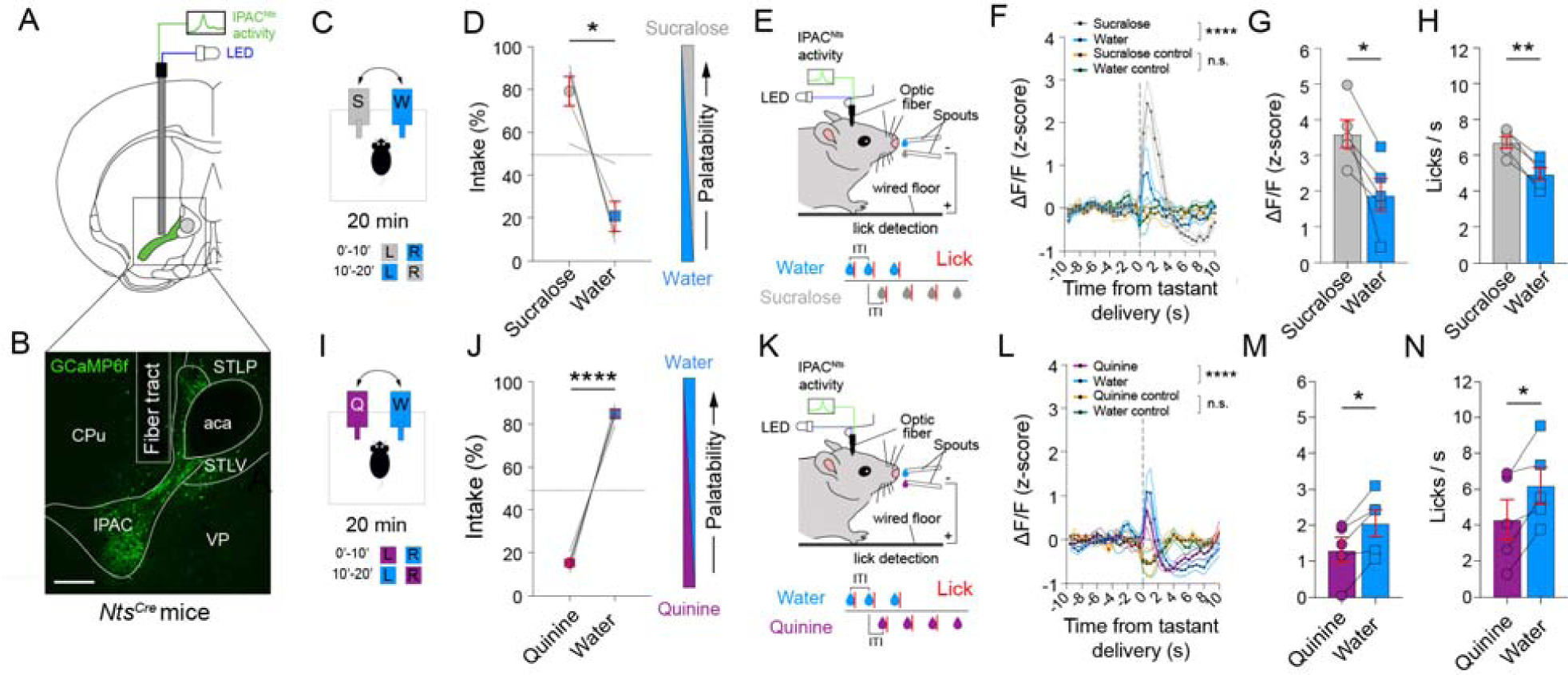
IPAC^Nts^ neurons encode the palatability of tastants. (A) A schematic of the approach. (B) A confocal image showing GCaMP6 expression in IPAC^Nts^ neurons and an optical-fiber tract in a representative mouse. Scale bar 200 μm. (C) A schematic of the design of the 2-bottle preference test. L, left bottle, R, right bottle. (D) Quantification of the intake of sucralose or water relative to total fluid intake (n = 5 mice, *p = 0.0128, paired t-test). (E) A schematic of the experimental design. (F) GCaMP6 and control (isosbestic) signals from IPAC^Nts^ neurons in mice (n = 5) consuming sucralose solution or water. GCaMP6 signals, F(39, 156) = 8.45, ****p < 0.0001; control signals, F(39, 156) = 1.19; p = 0.2278 (n.s.); two-way repeated-measures (RM) ANOVA, liquid x epoch interaction. (G) Peak GCaMP6 signals from IPAC^Nts^ neurons after the delivery of sucralose solution or water (n = 5 mice, *p = 0.0171, paired t-test). (H) Licking behavior after the delivery of sucralose solution or water, measured in a 3-s window following the first lick (n = 5 mice, **p = 0.0019, paired t-test). (I) A schematic of the design of the 2-bottle preference test. L, left bottle, R, right bottle. (J) Quantification of the intake of quinine or water relative to total fluid intake (n = 5 mice, ****p < 0.0001, paired t-test). (K) A schematic of the experimental design. (L) GCaMP6 and control (isosbestic) signals from IPAC^Nts^ neurons in mice (n = 5) consuming quinine solution or water. GCaMP6 signals, F(39, 156) = 4.054; ****p < 0.0001; control signals, F(39, 156) = 0.7166; p = 0.8883 (n.s.); two-way RM ANOVA, liquid x epoch interaction. (M) Peak GCaMP6 signals from IPAC^Nts^ neurons after the delivery of quinine solution or water (n = 5 mice, *p = 0.0145, paired t-test). (N) Licking behavior after the delivery of quinine solution or water (n = 5 mice, *p = 0.0218, paired t-test). Data are presented as mean ± s.e.m.

In human subjects, a common experimental approach to infer palatability is to present individuals with two foods with similar nutritional value (i.e., isocaloric) but different flavors. The difference in food intake is used as a measure of palatability (Yeomans, 1998; Yeomans and Wright, 1991). Using a similar approach, we tested the mice (which were under water restriction; Methods) for their preference for non-caloric solutions – sucralose, water and quinine – using the amount of intake as a proxy of tastant preference and thus palatability. These mice displayed a preference for sucralose over water (Figure 2C, D) and for water over quinine (Figure 2I, J), suggesting that sucralose and quinine are the most and least palatable tastants, respectively. Of note, as all these liquids are non-caloric, the preference should not be influenced by nutritional content.

*In vivo* fiber photometry revealed that IPAC^Nts^ neurons were robustly activated following liquid consumption (Figure 2E-H, K-N; Figure S2B-D). Notably, the activation by sucralose was greater than that by water (Figure 2F, G), and the activation by water was greater than that by quinine (Figure 2L, M). This ranking of neuronal activation mirrored the tastant preference measured behaviorally (Figure 2D, J). Thus, IPAC^Nts^ neuron activity scales with stimulus palatability. Analysis of the licking behavior showed that mice licked more vigorously at the spout delivering the preferred tastants (Figure 2 H, N), raising the possibility that IPAC^Nts^ activity could represent the motion associated with licking. However, we found no correlation between the amplitude of the neural response and lick rate in any of the mice (Figure S2E, F). Together, these results suggest that IPAC^Nts^ neuron activities represent tastant palatability, rather than the motor functions underlying licking.

### IPAC^Nts^ neurons integrate food stimulus information across sensory modalities

Besides the taste, the smell of a food has a major impact on its palatability (Yeomans, 1998). In mice, olfaction regulates feeding and metabolism (Patel et al., 2019; Riera et al., 2017). Appetitive food-related smells prepare inner organs for a meal (Brandt et al., 2018) and modulate the activity of homeostatic hypothalamic neurons (Chen et al., 2015).

To examine how IPAC^Nts^ neuron activities might be influenced by food-related smells, we used fiber photometry, as described above (Figure 2A, B; Figure S3A), to measure the *in vivo* responses of these neurons in food-restricted mice to the presentation of odors derived from different substances: a high-fat diet (HFD), butyric acid (BA), and mineral oil (MO, used to dissolve odorants and serving as a vehicle control; Methods; Figure 3A). While HFD is appetitive, BA is typically found in spoiled food and responsible for its rotten smell, and is thus aversive (Patel et al., 2019). Notably, we found that IPAC^Nts^ neurons were strongly activated by the smell of HFD, but were only minimally activated by the smell of BA or MO alone (Figure 3B-D). These results suggest that IPAC^Nts^ neurons are tuned to multiple features of palatable food, including both the taste and the smell.

**Figure 3.**
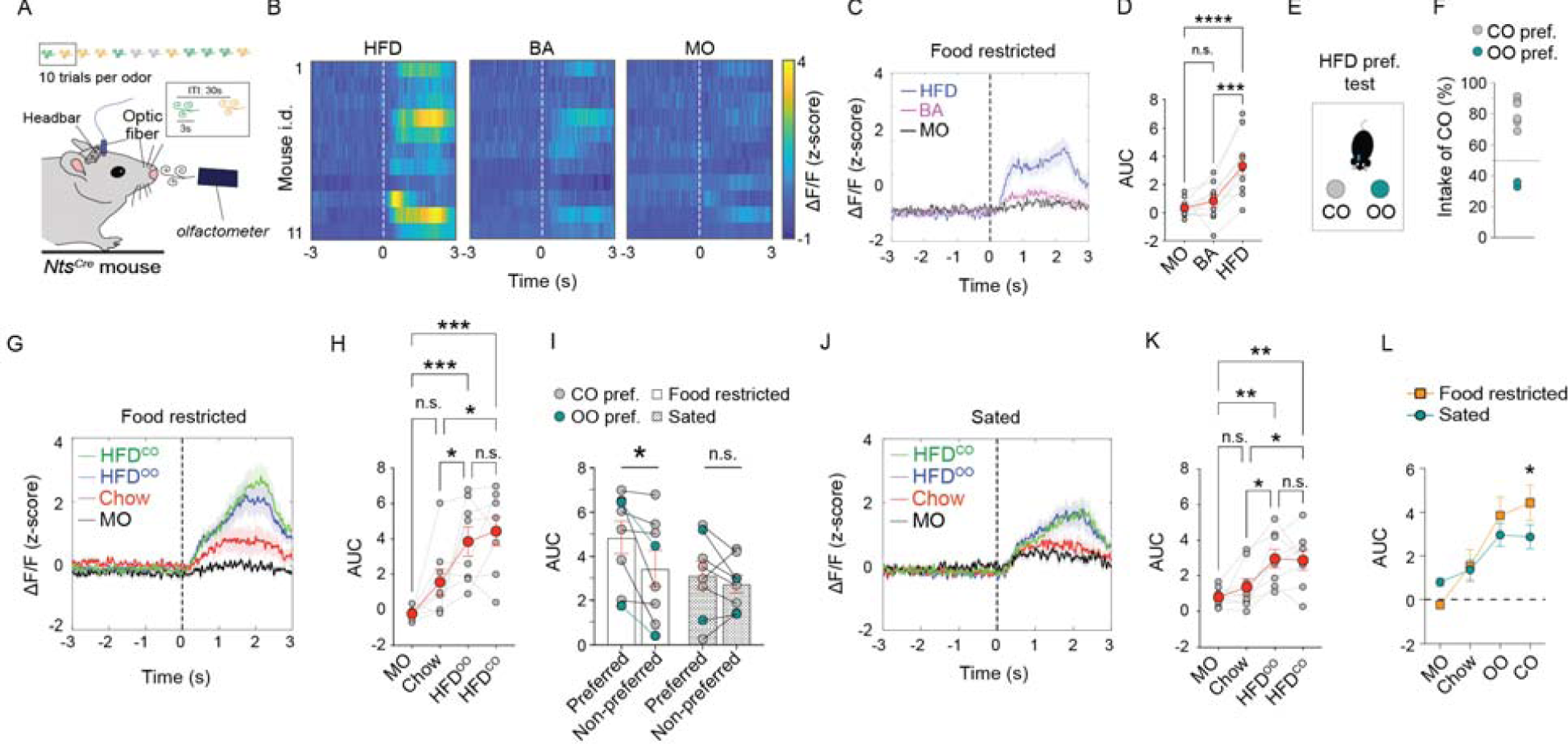
IPAC^Nts^ neurons encode the palatability of food odors. (A) A schematic of the experimental setup. (B) Heatmaps of average GCaMP6 responses of IPAC^Nts^ neurons in individual mice aligned to odor presentation (dashed line). HFD, high fat diet; BA, butyric acid; MO, mineral oil. (C) Average GCaMP6 signals from IPAC^Nts^ neurons in food-restricted mice aligned to HFD, BA and MO odor presentation (dashed line). (D) Quantification of the area under the curve (AUC) of the responses in individual mice between 0 and 3 s. N = 11 mice, F_(2,18)_ = 21.06, p < 0.0001; ***p < 0.001, ****p < 0.0001; One-way RM ANOVA followed by Holm-Sidak’s multiple comparisons test. (E) A schematic of the preference test. (F) Mice’s intake of HFD^CO^ relative to the total intake of HFD^CO^ and HFD^OO^. (G) Average GCaMP6 signals from IPAC^Nts^ neurons in food-restricted mice aligned to the presentation of different odors (dashed line). (H) Quantification of the AUC of the responses in individual mice between 0 and 3 s. N=8 mice, F_(3,21)_ = 11.96, p < 0.0001; n.s., p > 0.05, *p < 0.05, ***p < 0.001; one-way RM ANOVA followed by Holm-Sidak’s multiple comparisons test. (I) IPAC^Nts^ neurons responded more to the preferred HFD than the non-preferred in food-restricted mice (left), but not sated mice (right). N = 8 mice, F_(1,7)_ = 8.769, p = 0.0211; n.s., p > 0.05, *p < 0.05; two-way RM ANOVA followed by Holm-Sidak’s test. (J) Average GCaMP6 signals from IPAC^Nts^ neurons in sated mice aligned to the presentation of different odors (dashed line). (K) Quantification of the AUC of the responses in individual mice between 0 and 3 s. N=8 mice, F_(3,21)_ = 8.546, p = 0.0007; n.s., p > 0.05, *p < 0.05, **p < 0.01; one-way RM ANOVA followed by Holm-Sidak’s multiple comparisons test. (L) Average responses of IPAC^Nts^ neurons in (H) and (K) are replotted for visual inspection. F_(3,21)_ = 5.394, p = 0.0065; *p < 0.05, two-way RM ANOVA followed by Holm-Sidak’s multiple comparisons test. Data are presented as mean ± s.e.m.

### IPAC^Nts^ neurons represent the palatability of naturalistic food stimuli

In humans, the preference for a food, or food palatability, tends to be idiosyncratic. For instance, while some people prefer sweet food over spicy, others may do just the opposite. To assess the palatability of naturalistic foods in mice, we presented them with two kinds of HFDs that had identical nutritional value and macronutrient composition, but differed in lipid content, with one derived from coconut oil (HFD^CO^) and the other from olive oil (HFD^OO^). Mice avidly consumed either the HFD^CO^ or the HFD^OO^ pellets, even when sated (data not shown), indicating that both are highly rewarding and more palatable than chow, which was available *ad libitum* in the home cage. Interestingly, when both HFDs were offered, mice showed a clear preference for one of them, with the majority favoring HFD^CO^ over HFD^OO^ (Figure 3E, F). Since these diets were isocaloric, this preference should be dependent only on their palatability, but not the nutritional value.

To determine whether IPAC^Nts^ neuron activities encode food palatability, we presented food-restricted mice with odors derived from HFD^CO^, HFD^OO^, regular chow and MO. We found that IPAC^Nts^ neurons showed higher responses to the odors from the HFDs (HFD^CO^ and HFD^OO^) than the odor from regular chow or MO (Figure 3G, H; Figure S3B), paralleling the observation that all mice preferred HFDs over regular chow. Notably, when the idiosyncratic preferences of individual mice for one of the two HFDs (Figure 3E, F) were considered, IPAC^Nts^ neuron responses were larger for the odor of the favorite diet, irrespective of it being CO- or CO-flavored (Figure 3I, left).

In a separate experiment, we examined the responses of IPAC^Nts^ neurons in food-restricted mice to the smells of white chocolate (WCh) and dark chocolate (DCh) – which are isocaloric and among the popular energy-dense “cafeteria foods” causing obesity in humans – as well as regular chow and MO. We found that IPAC^Nts^ neurons preferentially responded to the smells of chocolates, especially WCh (Figure S3C-E). These results, together with those from the cFos experiments (Figure 1), strongly support the notion that IPAC^Nts^ neurons encode food palatability or preference.

Palatable food’s sensory attributes are sufficient to override homeostatic regulation, thereby driving excessive caloric intake (Morales and Berridge, 2020). To test whether IPAC^Nts^ neurons would respond to palatable food cues even in the absence of homeostatic drive, we repeated the above experiments in sated mice. We found that IPAC^Nts^ neurons still responded to food smells, and the magnitude of the response was larger for energy-dense foods (HFD^CO^, HFD^OO^, WCh, and DCh) than for chow or MO (Figure 3J, K; Figure S3B, C, F, G). Thus, IPAC^Nts^ neurons have preferential responses to the odors of palatable foods, even in sated mice.

Hunger is known to increase the palatability of a food, a phenomenon called alliesthesia (Cabanac, 1971). If IPAC^Nts^ neuron activity represents palatability, then such activity should be modulated by animal’s homeostatic state. Indeed, although IPAC^Nts^ neurons in hungry mice showed increased response to the favored HFD, this increase disappeared in sated mice (Figure 3I, right). Moreover, the response of IPAC^Nts^ neurons – especially that to the HFD – was larger when mice were in a hungry state than in satiety (Figure 3L). These results suggest that IPAC^Nts^ neuron activity is modulated by both food palatability and the homeostatic state of the animal. Together, our observations thus far unravel that IPAC^Nts^ neurons encode the palatability of both simple tastants and complex, real-world food stimuli, and point to the possibility that these neurons might have a role in regulating hedonic feeding, i.e., feeding in the absence of an energy deficit.

### Activation of IPAC^Nts^ neurons preferentially increases the intake of palatable foods

To assess whether activation of IPAC^Nts^ neurons could result in overfeeding, we selectively activated these neurons in sated mice with optogenetics. For this purpose, we bilaterally injected the IPAC of *Nts^Cre^* mice with an AAV expressing the light-gated cation channel channelrhodopsin (ChR2), or GFP (as a control), in a *Cre*-dependent manner. Optical fibres were implanted over the infected areas for light delivery (Figure 4A, Figure S4A). Following recovery from surgery and viral expression, sated mice received pellets of differing palatability (Figure 4B; Methods). During each food presentation, pulses of blue light were delivered into the IPAC.

**Figure 4.**
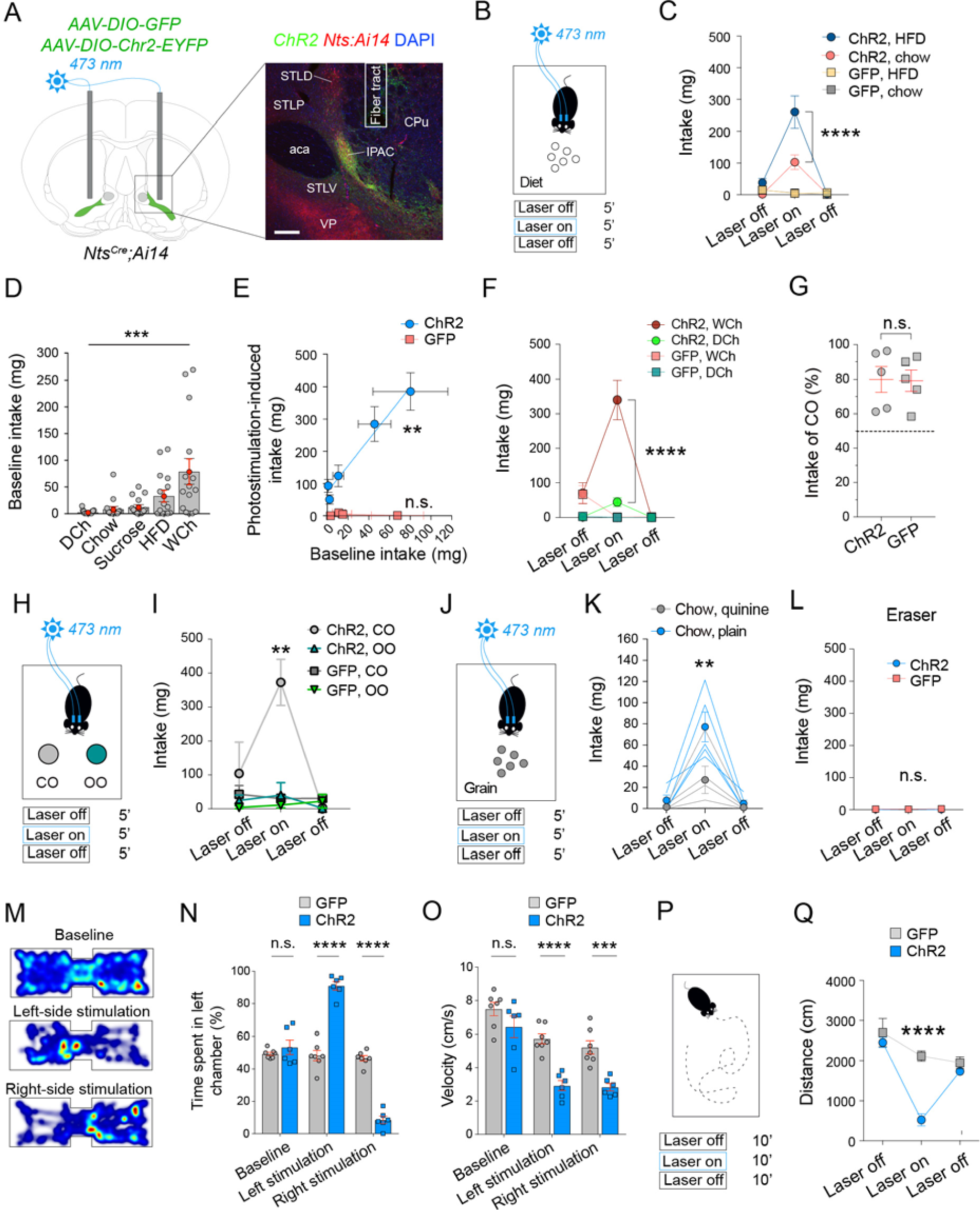
Activation of IPAC^Nts^ neurons selectively increases feeding on palatable foods. (A) Left: a schematic of the approach. Right: a confocal image showing ChR2 expression in IPAC^Nts^ neurons and an optical-fiber tract in a representative mouse. Scale bar 200 μm. (B) A schematic of the paradigm for testing the effects of optogenetics on feeding behavior. (C) Light delivery into the IPAC preferentially increased the intake of HFD over chow in the ChR2, but not GFP mice. ChR2 mice (n = 9): F_(2,16)_ = 12.64, p = 0.0005, ****p < 0.0001; GFP mice (n = 8): F_(2,14)_ = 0.6838, p = 0.5208; two-way RM ANOVA followed by Sidak’s multiple comparison test. (D) Food intake of sated mice (n = 15) at baseline (laser-off period) (***p = 0.0003, Friedman RM test). (E) Analysis of the relationship between food intake at baseline and during photostimulation. ChR2 mice: n = 9, **p = 0.0018, Pearson’s test; GFP mice: n = 6-8, p > 0.05 (n.s.), Spearman’s test. (F) Effect of light delivery into the IPAC on chocolate consumption. ChR2 mice (n = 9): F_(2,16)_ = 19.12, p < 0.0001, ****p < 0.0001; GFP mice (n = 6): F_(2,10)_ = 5.6, p = 0.0234; between WCh and DCh during laser on, p > 0.05; two-way RM ANOVA followed by Sidak’s multiple comparisons test. (G) Mice’s intake of HFD^CO^ relative to the total intake of HFD^CO^ and HFD^OO^ (n = 5 mice per group, p = 0.9385 (n.s.), unpaired t-test). (H) A schematic of the paradigm for testing the effect of optogenetics on food preference in sated mice. (I) Effect of light delivery into the IPAC on the consumption of HFD^CO^ or HFD^OO^. ChR2 mice (n = 5): F_(2,8)_ = 9.443, p = 0.0078, **p < 0.01; GFP mice (n = 5): F_(2,8)_ = 0.9049, p = 0.4423; two-way RM ANOVA followed by Sidak’s multiple comparisons test. (J) A schematic of the paradigm for testing the effect of optogenetics on food consumption in sated mice. (K) Effect of light delivery into the IPAC of the ChR2 mice on the consumption of quinine-flavored chow or plain chow (n = 5): F_(2,8)_ = 9.476, p = 0.0078, **p < 0.01, two-way RM ANOVA followed by Sidak’s multiple comparisons test. (L) Effect of light delivery into the IPAC of the ChR2 (n = 9) or GFP (n = 6) mice on the consumption of inedible items (i.e., pencil eraser) (F_(2,26)_ = 1.066, p = 0.3591 (n.s.), two-way RM ANOVA). (M) Heatmaps for the activity of a representative mouse at baseline (top), or in a situation whereby entering the left (middle) or right (bottom) side of the chamber triggered photostimulation in the IPAC. (N) Preference of ChR2 (n = 6) and GFP mice (n = 7) for the left chamber side (F_(2,22)_ = 137.9; p < 0.0001; ****p < 0.0001; two-way RM ANOVA followed by Sidak’s multiple comparisons test). (O) Velocity of the ChR2 (n = 6) and GFP (n = 7) mice in the RTPP/A task (F_(2,24)_ = 7.116, p = 0.0041; ***p < 0.001; ****p < 0.0001; two-way RM ANOVA followed by Sidak’s multiple comparisons test). (P) A schematic of the open field test. (Q) Distance traveled during the open field test for the ChR2 (n = 8) and GFP (n = 6) mice (F_(2,24)_ = 13.37, p < 0.0001; ****p < 0.0001; two-way RM ANOVA followed by Sidak’s multiple comparisons test). Data are presented as mean ± s.e.m.

We found that photostimulation in the ChR2 mice increased their intake of both regular chow and the more palatable HFD, and, interestingly, the effect was larger for HFD than regular chow (Figure 4C). Consistently, the number and length of feeding bouts were also increased by the photostimulation (Figure S4B-D). Photostimulation in these mice occasionally resulted in stereotyped appetitive behaviors such as licking the floor, foraging, and gnawing of inedible bedding pellets (Supplementary Video 1 & 2). In contrast, photostimulation in the GFP mice had no behavioral effect (Figure 4C; Figure S4C-E). These results indicate that activation of IPAC^Nts^ neurons drives food intake in the absence of metabolic need and even more so for palatable food.

These data, together with the photometry results, led us to hypothesize that IPAC^Nts^ neuron activity promotes feeding as a function of food palatability. To test this hypothesis, we activated IPAC^Nts^ neurons with optogenetics in sated mice, as described above (Figure 4B, C), and assessed the effects of this manipulation on the consumption of chow, HFD, WCh, DCh, and sucrose. These foods have different palatability, as indicated by mice’s differential preference for them before the activation (baseline intake, Figure 4D). We found that activation of IPAC^Nts^ neurons promoted the intake of all these foods (Figure 4E; S4E), and, remarkably, there was a strong correlation between the activation-induced intake and baseline intake (Figure 4E). These results suggest that the feeding-promoting effect of IPAC^Nts^ neuron activation is indeed dependent on palatability.

As these foods have different nutritional values, which may influence consumption independent of palatability, we repeated the above experiments with three pairs of foods that are isocaloric but differ in palatability: (1) WCh and DCh (Figure 4F), (2) HFD^CO^ and HFD^OO^ (Figure 4G-I), and (3) plain chow and chow flavored with quinine (Figure 4J, K; Methods). WCh was much more preferred than DCh by all mice, likely due to the bitter taste of the latter (Figure 4D). The preference for HFD^CO^ or HFD^OO^ was more idiosyncratic (see Figure 3E, F), but all mice in this group preferred HFD^CO^ (Figure 4G). For the plain and quinine-flavored chow, the two kinds of food pellets had identical texture and nutritional value, but the quinine-flavored was expected to be less palatable (Nisbett, 1968).

We found that in all three cases, activation of IPAC^Nts^ neurons dramatically increased the intake of the more palatable food and increased the intake of the less palatable counterpart in the pair to a much lesser degree (Figure 4F-K). Of note, IPAC^Nts^ activation did not induce any feeding on a random inedible object, such as an eraser (Figure 4L), suggesting the effect is food-specific. Together, these results strongly indicate that IPAC^Nts^ neuron activity promotes food intake in a palatability-dependent manner, and thus may play an important role in controlling hedonic feeding.

### Activation of IPAC^Nts^ neurons is positively reinforcing and decreases locomotor activity

Previous studies indicate that activating a BNST-to-LHA circuit, which regulates metabolic feeding, is rewarding and can support intracranial self-stimulation (Jennings et al., 2013). Interestingly, this phenomenon was dependent on the homeostatic state of the animal, as satiety and food-restriction significantly decreased and increased, respectively, the degree of self-stimulation (Jennings et al., 2013). To determine whether IPAC^Nts^ neurons act in a similar manner, we presented the ChR2 and GFP mice – in which the IPAC^Nts^ neurons expressed ChR2 and GFP, respectively – with two ports (Figure S4F). Poking into one of the ports (the active port) would lead to the delivery of light pulses into the IPAC, whereas poking into the other port (the inactive port) would have no consequence. The ChR2 but not GFP mice consistently poked into the active port to receive the photostimulation while completely ignoring the inactive port, demonstrating that activation of IPAC^Nts^ neurons effectively supports self-stimulation (Figure S4G, H). Interestingly, there was no difference in self-stimulation rates when the mice were tested sated on regular chow or a high fat diet, or following food-restriction (FR) (Figure S4I, J; Methods). These results indicate that activation of IPAC^Nts^ neurons is intrinsically rewarding. Moreover, the rewarding effect is independent of animal’s homeostatic state, and thus different from the function of the circuits regulating metabolic feeding (Jennings et al., 2013).

In line with the self-stimulation results, photo-activation of IPAC^Nts^ neurons induced place preference in a real-time place preference or aversion (RTPP/A) assay (Figure 4M, N; Methods). Notably, we found that activation of IPAC^Nts^ neurons caused a significant reduction in mice’s movements in both the RTPP/A assay (Figure 4O) and an open field test (Figure 4P, Q; Methods). These results show that IPAC^Nts^ neuron activation is positively reinforcing, and moreover suggest that these neurons may regulate animal’s physical activity, a major channel for energy expenditure, in addition to influencing energy intake.

### Inhibition of IPAC^Nts^ neurons promotes energy expenditure

To determine whether the activity of IPAC^Nts^ neurons is required for energy homeostasis, we selectively blocked neurotransmitter release from these neurons with the tetanus toxin light chain (TeLC) (Murray et al., 2011). For this purpose, we bilaterally injected the IPAC of *Nts^Cre^* mice with an AAV expressing TeLC, or GFP (as a control), in a Cre-dependent manner (Figure 5A, B). Mice were returned to their home cage after the surgery. Although the TeLC mice and GFP mice had similar body weight prior to the surgery (Figure S5A; “day 0” (d0) timepoint), the TeLC group dramatically lost weight within 10 days post-surgery and did not regain it by d30 (Figure S5A; Figure 5C). On average, at d30, the GFP mice gained 4.0 ± 2.0% of their initial body weight, whereas the TeLC mice lost 17.2 ± 4.3% (Figure S5A). Notably, the sudden loss of mass in the TeLC mice did not lead to starvation, as their body weight stabilized between d10 and d30 (Figure S5A, right).

**Figure 5.**
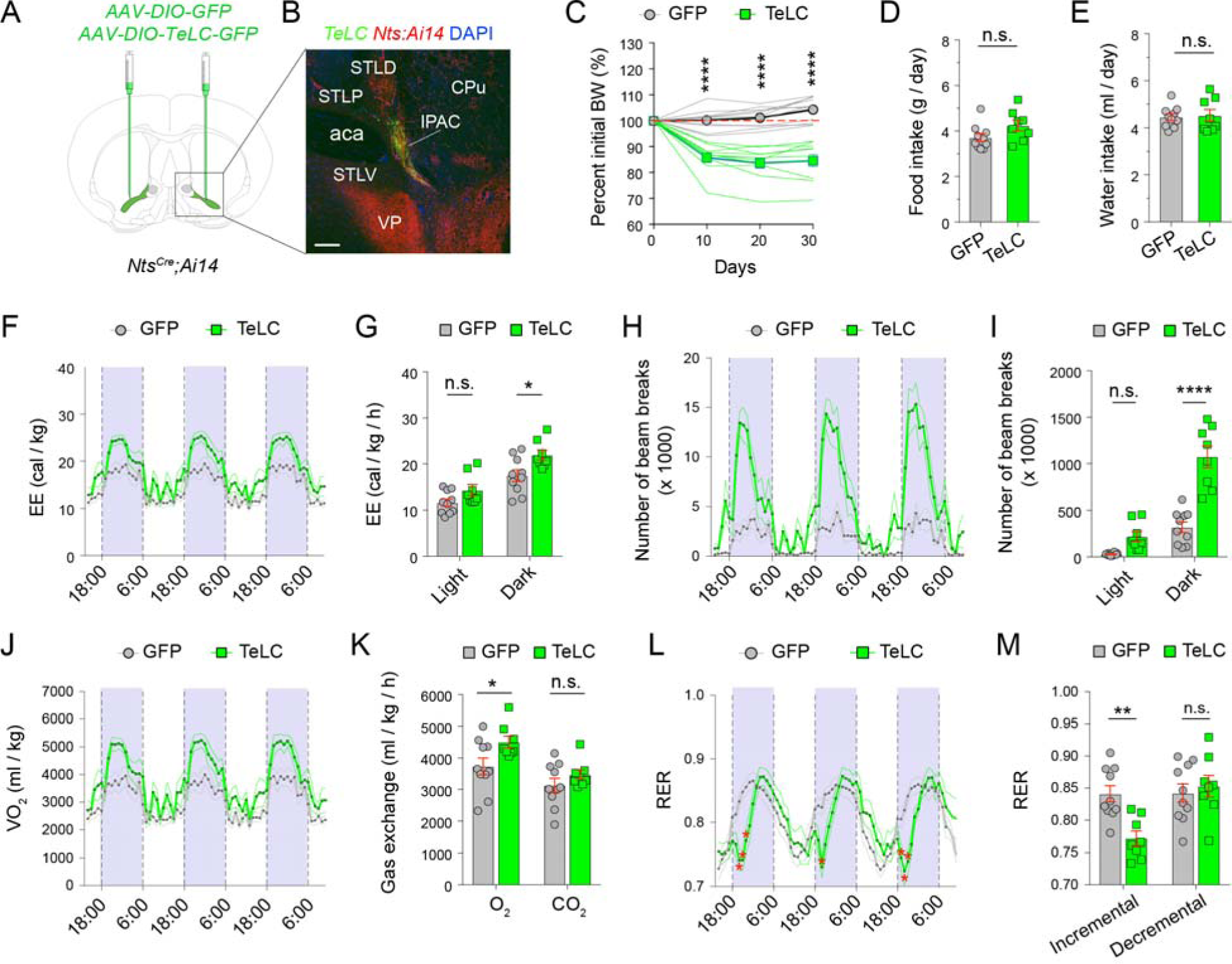
Inhibition of IPAC^Nts^ neurons increases energy expenditure. (A) A schematic of the approach. (B) A confocal image showing TeLC expression in IPAC^Nts^ neurons in a representative mouse. Scale bar 200 μm. (C) Changes in bodyweight (BW) in the GFP mice (n = 11) and TeLC mice (n = 10) following viral injection (d0) (F_(3,57)_ = 44.28, p < 0.0001; ****p < 0.0001; two-way RM ANOVA followed by Sidak’s multiple comparisons test). (D) Daily food (chow) intake of the GFP mice (n = 10) and TeLC mice (n = 8) (p = 0.0785 (n.s.), unpaired t-test). (E) Daily water intake of the GFP mice (n = 10) and TeLC mice (n = 8) (p = 0.8023 (n.s.), unpaired t-test). (F) Energy expenditure of the GFP mice (n = 10) and TeLC mice (n = 8) over 72 h. Data are plotted in 1-h intervals. White and purple areas represent light (6:00-18:00) and dark cycles (18:00-6:00), respectively (F_(70, 1120)_ = 2.029, p < 0.0001, two-way RM ANOVA). (G) Average energy expenditure of the mice in (F) during light and dark cycles (F_(1, 16)_ = 5.934, p = 0.0269; *p < 0.05, n.s., p > 0.05; two-way RM ANOVA followed by Sidak’s multiple comparisons test). (H) Locomotor activity of the GFP (n = 10) and TeLC mice (n = 8) over 72 h (F_(70,1120)_ = 7.699, p < 0.0001, two-way RM ANOVA). (I) Average locomotor activity of the mice in (H) during light and dark cycles (F_(1,16)_ = 37.84, p < 0.0001; ****p < 0.0001, n.s., p > 0.05, two-way RM ANOVA followed by Sidak’s multiple comparisons test). (J) The volume of oxygen consumed (VO_2_) by GFP (n = 10) and TeLC mice (n = 8) over 72 h (F_(70,1120)_ = 2.221, p < 0.0001, two-way RM ANOVA). (K) O_2_ and carbon dioxide (CO_2_) exchange during incremental activities in the dark cycle (18:00-6:00) (GFP mice, n = 10, TeLC mice, n = 8; F_(1,16)_ = 20.24, p = 0.0004; *p < 0.05, n.s., p > 0.05; two-way RM ANOVA followed by Sidak’s multiple comparisons test). (L) Respiratory exchange ratio (RER) of GFP (n = 10) and TeLC mice (n = 8) over 72 h (F_(70,1120)_ = 5.042, p < 0.0001, two-way RM ANOVA). (M) Average RER during incremental and decremental activities in the dark cycle (18:00-6:00) on the 3^rd^ day for the mice in (L) (F_(1,16)_ = 20.24, p = 0.0005; **p < 0.01, n.s., p > 0.05, two-way RM ANOVA followed by Sidak’s multiple comparisons test). The effect was similar in other days.

As a reduction in body weight is normally caused by an imbalance between energy intake and expenditure, we next examined the source of the imbalance in these mice by monitoring their food (and water) intake and energy expenditure for 72 h in metabolic cages (Methods). Prior to the examination, the TeLC and GFP mice lost 15.1±2.1% and gained 1.7±1.6 %, respectively, of their initial body weight (Figure S5B). No further change in body weight was observed in either cohort within this 72-h period (Figure S5C), suggesting that the TeLC mice reached a new energy balance after the initial weight loss. In addition, we found no difference in the gross food or water intake in the TeLC mice compared to controls (Figure 5D, E), suggesting that the neural circuits underlying hunger and thirst homeostasis were minimally affected by inhibition of IPAC^Nts^ neurons. Interestingly, energy expenditure (EE) was significantly higher in the TeLC mice than controls, in particular during the active phase (i.e., dark cycle) of the day for mice (Figure 5F, G).

Energy is expended to sustain basal metabolic rate, the thermic effect of food and physical activities. We reasoned that an increase in physical activity accounts for the increase in energy expenditure in the TeLC mice, because mice with enhanced IPAC^Nts^ neuron activity – opposite to the effect of TeLC – showed decreased locomotor activity (Figure 4P, Q). Indeed, we found that the TeLC mice were 3.7-fold more active than controls (Figure 5H, I; Figure S5D). In humans, the maximum volume of oxygen inhaled (VO_2_^Max^) and the respiratory exchange ratio (RER, the ratio between VCO_2_ produced and VO_2_ inhaled) during exercise are gold standards for endurance and cardiovascular fitness (Gollnick, 1985; Ramos-Jimenez et al., 2008). Athletes, for example, typically have higher VO ^Max^ and lower RER than untrained subjects during incremental exercise (which increases in intensity over time until VO_2_^Max^ is reached), with the lower RER indicative of a higher lipid oxidation (i.e., fat burning) rate. Notably, we found that the TeLC group had markedly higher VO_2_ and lower RER than the control group during the period in a day when mice had “incremental activities” (from the beginning of the dark phase to when the highest VO_2_ levels were reached) (Figure 5J-M; Figure S5E-G). These results suggest that the TeLC mice were physically fitter than the controls.

To determine whether our manipulation affected anxiety-related behaviors, we subjected the TeLC mice and GFP control mice to the elevated-plus maze (EPM) and open-field (OF) tests (Figure S6). We found no difference between the two groups in measures of anxiety behaviors in rodents, including the time spent on the open arms of the EPM (Figure S6A, B) or in the center of the OF (Figure S6E, F), or the frequency of entering those spaces (Figure S6C, G). Interestingly, the two groups also had similar locomotor activities during these anxiogenic tests (Figure S6D, H), suggesting that the increase in physical activities in the TeLC mice is context- or state-dependent, and may reflect volitional activities in a familiar or less stressful environment.

Together, these results show that inhibition of IPAC^Nts^ neurons dramatically increases energy expenditure, aerobic capacity, and physical fitness without affecting regular food intake, suggesting that the activity of these neurons normally constrains energy expenditure by limiting physical activity. However, the activity of these neurons is not critical for homeostatic energy intake.

### Inhibition of IPAC^Nts^ neurons decreases food palatability

Our photometry and optogenetic activation results indicate that IPAC^Nts^ neuron activity encodes food palatability and is sufficient to promote hedonic feeding. We thus tested whether the activity of these neurons is also required for this type of feeding. To this end, we switched the diet from regular chow to HFD for the mice and subsequently assessed their food and water intake (Figure 6A). We found that, in the four days following the switch, the GFP mice ate significantly more than the TeLC mice (Figure 6B), however this was not observed when the mice were fed with chow (Figure 5D). Accordingly, water intake was lower for TeLC mice on HFD (Figure 6C). To assess the change in energy intake (EI) after the diet switch, we calculated the daily calorie consumption when fed with chow (Figure 5D) or HFD (Figure 6B) and found that the GFP, but not TeLC mice, dramatically increased EI when fed with HFD (Figure 6D). EI increased by 40% in the GFP mice but remained unchanged in the TeLC mice (Figure 6E), suggesting that the GFP, but not TeLC mice, ate above their metabolic needs and likely entered a positive energy balance state. Indeed, after only 4 days of HFD, the body weight of GFP mice dramatically increased while that of TeLC mice remained stable (Figure 6F). These data strongly suggest that inhibition of IPAC^Nts^ neurons prevents overfeeding on palatable foods and acute weight gain.

**Figure 6.**
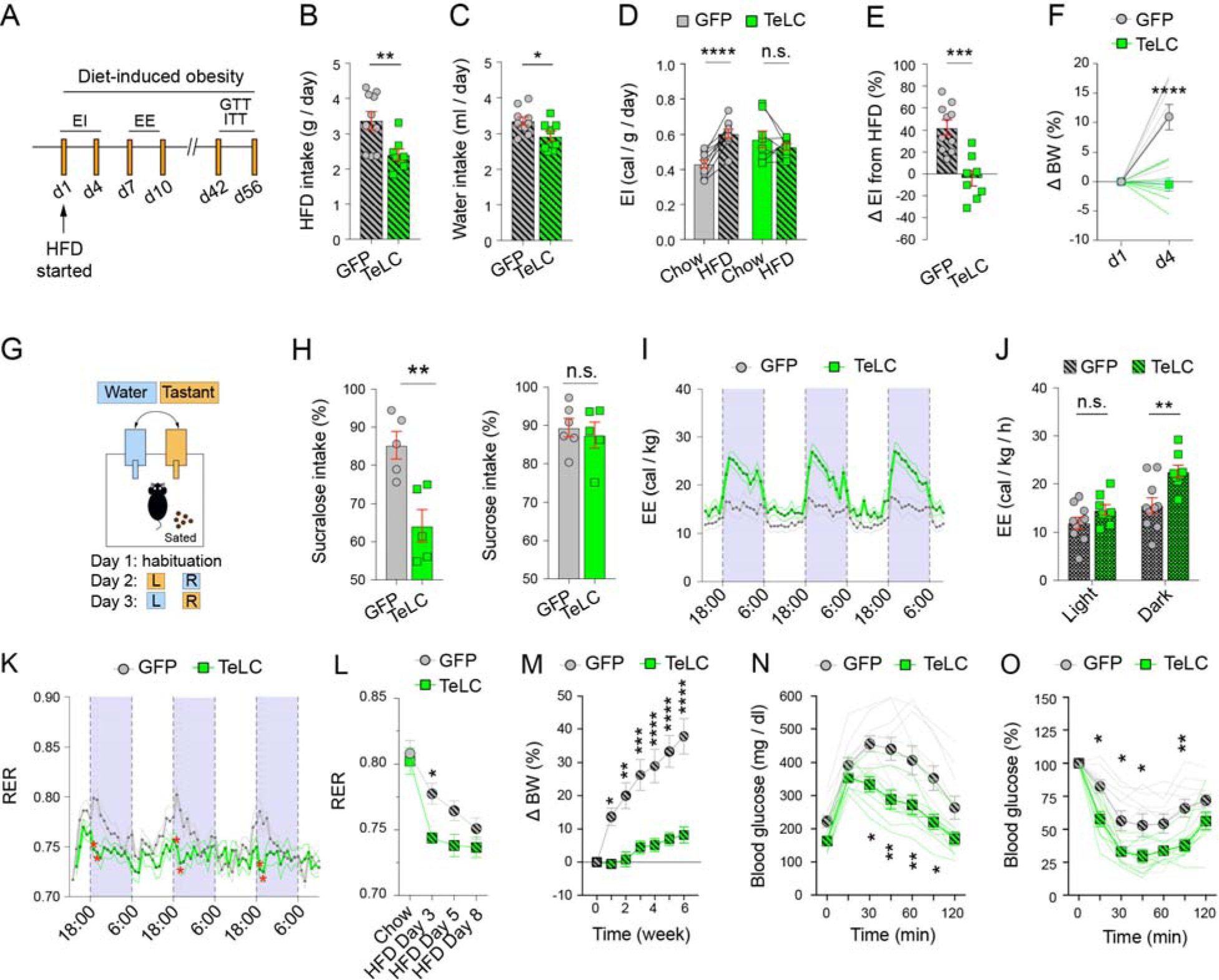
Inhibition of IPAC^Nts^ neurons protects from HFD-induced weight gain. (A) A schematic of the experimental design. GTT and ITT, glucose and insulin tolerance tests, respectively. (B) Daily HFD intake (GFP mice, n = 10, TeLC mice, n = 8; **P = 0.0073, unpaired t-test). (C) Daily water intake of the same mice in (B) (*P = 0.0305, unpaired t-test). (D) Comparison of energy intake from chow and HFD diets (derived from Figure 5D; GFP, n = 10, ****p < 0.0001; TeLC mice, n = 8, p = 0.3562 (n.s.); paired t-test). (E) Change in energy intake after the switch from chow to HFD, in the same mice as those in (D) (***P = 0.0002, unpaired t-test). (F) Acute changes in bodyweight (BW) following 4 days of HFD, in the same mice as those in (D) (F_(1,16)_ = 19.45, p = 0.0004, ****p < 0.0001, two-way RM ANOVA followed by Sidak’s multiple comparisons test). (G) A schematic of the design of the 2-bottle preference test. L, left bottle, R, right bottle. (H) Left: quantification of the intake of sucralose relative to total fluid intake (GFP mice, n = 5, TeLC mice, n = 5; **p = 0.0055, unpaired t-test). Right: quantification of the intake of sucrose relative to total fluid intake (GFP mice, n = 6, TeLC mice, n = 5; p = 0.6488 (n.s.), unpaired t-test). (I) Energy expenditure of the GFP mice (n = 10) and TeLC mice (n = 8) over 72 h, when fed HFD. Data are plotted in 1-h intervals. White and purple represent light (6:00-18:00) and dark cycles (18:00-6:00), respectively (F_(71, 1136)_ = 7.087, p < 0.0001, two-way RM ANOVA). (J) Average energy expenditure of the HFD-fed mice in (I) during light and dark cycles (F_(1, 16)_ = 6.527, p = 0.0212; **p < 0.01, n.s., p > 0.05; two-way RM ANOVA followed by Sidak’s multiple comparisons test). (K) RER values of the HFD-fed mice over 72 h (GFP mice, n = 10, TeLC mice, n = 8; F_(71, 1136)_ = 2.337, p < 0.0001; *p < 0.05; two-way RM ANOVA followed by Sidak’s post hoc multiple comparisons test). (L) Quantification of the changes in the RER (GFP mice, n = 10, TeLC mice, n = 8; F_(1, 16)_ = 10.71, p = 0.0048; *p < 0.05; two-way RM ANOVA followed by Sidak’s multiple comparisons test). (M) Change in bodyweight (BW) after switching to HFD diet (GFP mice, n = 10, TeLC mice, n = 8; F_(6, 120)_ = 16.9, p < 0.0001; *p < 0.05, **p < 0.01, ***p < 0.001, ****p < 0.0001; two-way RM ANOVA followed by Sidak’s multiple comparisons test). (N) Blood glucose levels following glucose administration during GTT (see A) (GFP mice, n = 10, TeLC mice, n = 8; F_(6, 96)_ = 4.37, p = 0.0006; *p < 0.05, **p < 0.01; two-way RM ANOVA followed by Sidak’s multiple comparisons test). (O) Blood glucose levels following insulin administration during ITT in the same mice as those in (N) (F_(6, 96)_ = 2.794, p = 0.0151; *p < 0.05, **p < 0.01; two-way RM ANOVA followed by Sidak’s multiple comparisons test).

To determine whether an impairment in detecting nutritional value contributed to the preventive effect on overfeeding, we tested these mice’s preference for a sucralose solution versus water, or a sucrose solution versus water (Figure 6G; Methods). Sucralose and sucrose were used because they are both sweet tastants, but are noncaloric and caloric, respectively. Notably, the TeLC mice showed decreased preference for sucralose, but normal preference for sucrose compared with the GFP control mice (Figure 6H). This result suggests that IPAC^Nts^ neuron activity is required for the orosensory perception of a palatable stimulus (i.e., palatability), but is dispensable for detecting the nutritional value of the stimulus.

### Inhibition of IPAC^Nts^ neurons prevents obesity and facilitates glucose metabolism

Following one week of feeding on HFD, TeLC mice still displayed markedly elevated energy expenditure (Figure 6I, J) and locomotor activity (Figure S7A, B), which were accompanied by increased VO_2_ and VCO_2_ (Figure S6C-F) and decreased RER (Figure 6K, L) compared with the GFP mice. As a result, the TeLC mice had lower body weight-gain than the GFP mice during this period (Figure 6M, first week).

Continued feeding on HFD (Figure 6A) effectively led to diet-induced obesity (DIO) in the GFP mice, as these mice steadily gained weight over the course of several weeks (Figure 6M). In stark contrast, the TeLC mice remained lean despite the HFD (Figure 6M), a phenotype that prompted us to investigate possible beneficial effects of IPAC^Nts^ neuron inhibition on glucose metabolism. We found that blood glucose levels were significantly lower in the TeLC mice than GFP mice in a glucose tolerance test (Figure 6N; Methods). Moreover, the TeLC mice showed lower glucose levels when measured in an insulin sensitivity test (Figure 6O; Methods). These results suggest that inhibition of IPAC^Nts^ neurons confers resistance to DIO and ameliorates its detrimental effects on glucose tolerance and insulin sensitivity.

The weight of the TeLC mice was 27.2% lower than the GFP mice at experiment endpoint (8 weeks of HFD; Figure 7A, B). Organ tissue analysis revealed no overt changes in the weight of different organs in the TeLC mice compared with the GFP mice (Figure 7C). However, the TeLC mice had 75.3% lower inguinal white adipose tissue (iWAT), 67.1% lower epididymal WAT (eWAT), 66.1% lower mesenteric WAT (mWAT) and 46.0% lower brown adipose tissue (BAT) (Figure 7D). Consistently, a lower amount of lipid droplets was found in the BAT (Figure 7E) and liver (Figure 7F) of the TeLC mice compared to controls. The adipocyte size in iWAT and eWAT was correspondingly decreased in these mice (Figure 7G, H). Interestingly, histological analysis revealed that the iWAT – but not the eWAT – of some TeLC mice contained multilocular lipid droplets (Figure 7G), suggesting lipid browning. In line with this observation, we found that the expression of *Ucp1* (*Uncoupling protein 1*) – which is usually highly expressed in BAT, but not WAT – was significantly higher in the iWAT of TeLC mice compared to controls (Figure 7I). Of note, *Ucp1* is critically involved in adaptive thermogenesis by uncoupling mitochondrial oxidative metabolism from ATP production (Cannon and Nedergaard, 2004).

**Figure 7.**
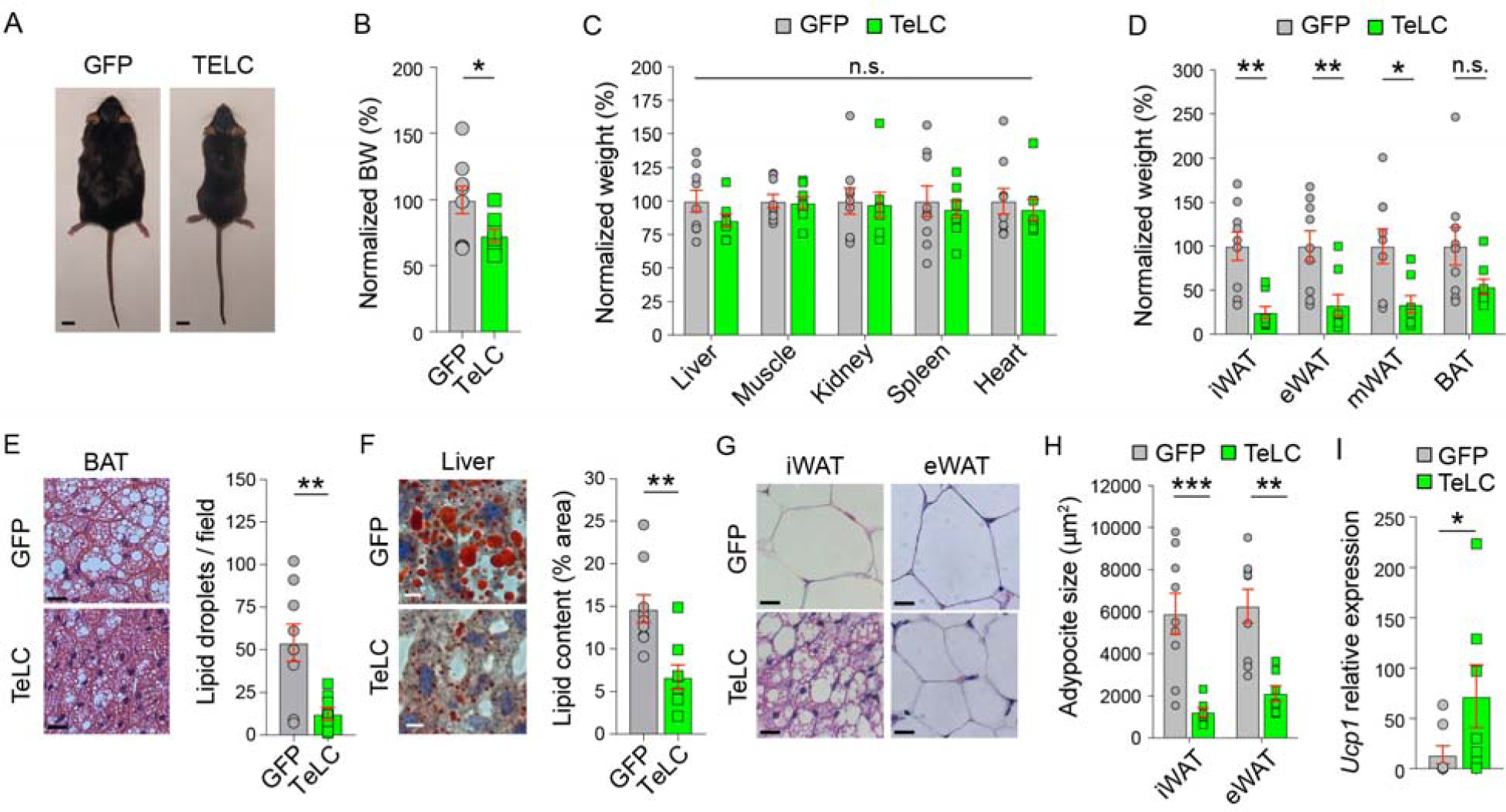
Inhibition of IPAC^Nts^ neurons reduces adiposity. (A) Representative images of a GFP and a TELC mouse at endpoint of the experiment (8 weeks of DIO). Scale bar: 1 cm. (B) Quantification of the bodyweight (BW) of the mice during DIO, which was normalized to the BW of GFP mice (GFP mice, n = 9, TeLC mice, n = 8; *p = 0.0381, unpaired t-test). (C) Quantification of the weight of different organs in the mice treated with DIO, with the weight of each organ normalized to that of GFP mice (GFP mice, n = 9, TeLC mice, n = 8; F_(1,15)_ =0.4306, n.s., p > 0.05, two-way ANOVA). (D) Quantification of the weight of different adipose tissues in the same mice as those in (C), with the weight normalized to that of GFP mice (F_(1,15)_ = 9.757, p = 0.0070; *p < 0.05, **p < 0.01, n.s., p > 0.05; two-way RM ANOVA followed by Sidak’s multiple comparisons test). (E) Left: representative images of BAT tissue stained for H&E from a GFP (top) and a TeLC (bottom) mouse treated with DIO. Scale bar: 20 μm. Right: quantification of the number of large lipid droplets in the two groups (GFP mice, n = 9; TeLC mice, n = 8; **p = 0.0037, unpaired t-test). (F) Left: representative images of liver tissue stained with Red-Oil from a GFP (top) and a TeLC (bottom) mouse treated with DIO. Scale bar: 10 μm. Right: quantification of the area occupied by lipid droplets in the two groups (GFP mice, n = 9, TeLC mice, n = 8; **p = 0.0025, unpaired t-test). (G) Representative images of WAT tissues stained for H&E from a GFP (top) and a TeLC (bottom) mouse treated with DIO. Scale bar: 20 μm. (H) Quantification of adipocyte size in iWAT and eWAT based on histological images as those in (G) (GFP mice, n = 8; TeLC mice, n = 7; F_(1,14)_ = 19.8, p = 0.0005; **p < 0.01, ***p < 0.001; two-way RM ANOVA followed by Sidak’s multiple comparisons test). (I) Expression of *Ucp1* in iWAT tissue from GFP and TELC mice treated with DIO (GFP mice, n = 8, TeLC mice, n = 7; *p = 0.0289, Mann Whitney U-test).

Together, these data show that inhibition of IPAC^Nts^ neurons protects from obesity by reducing caloric intake and increasing energy expenditure via elevated physical activity, thereby improving fitness and metabolic health indicators, such as VO_2_, RER, glucose metabolism and lipid browning.

## DISCUSSION

The easy availability of energy-dense foods is a main contributor to the current obesity pandemic. Foods rich in sugars and lipids are highly palatable, increasing hedonic intake (Morales and Berridge, 2020; Yeomans, 1998; Yeomans and Wright, 1991) that overrides homeostasis and leads to weight gain. Recent studies have identified neurons in different brain areas – such as the VP, NAc and peri-locus coeruleus – that encode or regulate hedonic feeding (Gong et al., 2020; Ottenheimer et al., 2018; Taha and Fields, 2005), but our understanding of the neural mechanisms underlying this behavior remains incomplete.

In this study, we show that IPAC^Nts^ neurons bidirectionally regulate palatable food intake and energy expenditure, thereby regulating peripheral organ function and energy homeostasis. In particular, these neurons respond to gustatory and olfactory food-related stimuli and encode their palatability. The magnitude of the response was correlated with food preference, modulated by the internal state of the animal, and independent of nutritional value. Furthermore, activation of IPAC^Nts^ neurons increased food consumption even in sated mice in a palatability-dependent manner, such that the effect was larger for more palatable foods and minimal for bitter foods or tastants. However, the effect did not depend on the caloric content of the foods. Thus, the activity of IPAC^Nts^ neurons likely contributes to dietary choice based on the hedonic, orosensory properties of foods.

Consistent with this notion, blockage of IPAC^Nts^ neurons reduced palatable food intake and food preference, but had no obvious effect on homeostatic feeding. Mice with IPAC^Nts^ neuron inhibition ate only the calories needed for sustaining their metabolic needs, while control mice overate on the HFD. Importantly, such effects of IPAC^Nts^ neuron inhibition seemed to be caused by a decreased ability to perceive the HFD orosensory properties, not by an impairment in detecting caloric compounds. These results are consistent with previous findings that separate neural substrates mediate the detection of hedonic and nutritional reward properties. For example, *Trpm5*-deficient mice are unable to sense sweet taste but are able to detect sucrose on the basis of its caloric content (Beeler et al., 2012; de Araujo et al., 2008); and specialized neurons in the NTS are known to encode the caloric value of glucose (Tan et al., 2020).

Lack of physical exercise, and thus energy expenditure, is another factor strongly contributing to obesity onset and progression (Hill et al., 2012; Tremblay and Willms, 2003). We found that activation of IPAC^Nts^ neurons caused a decrease in animals’ locomotor activity. Conversely, sustained inhibition of these neurons drastically increased locomotion, increased volume of oxygen uptake (VO_2_) and lowered Respiratory Exchange Ratio (RER). Both VO_2_ and RER are used to determine physical fitness in human subjects (Gollnick, 1985; Ramos-Jimenez et al., 2008). A lower RER indicates a higher rate of lipid oxidation (i.e., fat burning), thus providing an explanation for the decreased body weight following inhibition of IPAC^Nts^ neurons.

The combination of reduced HFD intake and increased energy expenditure in mice with IPAC^Nts^ neuron inhibition likely confers protection from the detrimental effects of chronic HFD feeding in the DIO model. Indeed, mice with impaired IPAC^Nts^ transmission remained lean and had improved glucose homeostasis compared with controls when challenged with DIO, suggesting that suppressing IPAC^Nts^ neuron activity effectively protects from metabolic diseases.

The reduced body weight of mice with IPAC^Nts^ neuron inhibition was mostly represented as a dramatic reduction in white adipose storage. Indeed, histological analysis in these mice revealed that iWAT and eWAT adipocytes were visibly smaller than those in control mice, and furthermore, exhibited features of BAT (e.g., multilocular appearance). Interestingly, IPAC^Nts^ neuron inhibition increased the expression of *Ucp1* in the iWAT, which is usually highly expressed in BAT (but not WAT) and regulates thermogenesis (Aldiss et al., 2018). Thus, inhibition of IPAC^Nts^ neurons induces changes in not only energy intake and expenditure, but also energy storage. Importantly, inhibition of IPAC^Nts^ neurons reduces lipid droplet accumulation in the BAT and liver in the DIO model, which is characteristic of BAT dysfunction (i.e., BAT “whitening”) (Shimizu et al., 2014) and nonalcoholic fatty liver disease (Recena Aydos et al., 2019) associated with obesity in humans.

Collectively, our results identify IPAC^Nts^ neurons as a crucial substrate for regulating energy balance-related behaviors, including hedonic food intake and energy expenditure/deposit homeostasis. In particular, given previous findings in humans that BAT is linked to metabolic health (Matsushita et al., 2014), and exercise is known to improve metabolic conditions and reduce the risk for or improve the prognosis of metabolic diseases (Cormie et al., 2017; Moore et al., 2016; Moore et al., 2012; Nayor et al., 2020), our results suggest that manipulation of IPAC^Nts^ neurons might have important implications in the prevention or treatment of these disorders, including bodyweight-related disorders and cancer.

## ACKNOWLEDGEMENTS

We thank Taylor Russo for technical assistance, and members of the Li laboratory for helpful discussions. This work was supported by grants from EMBO (ALTF 458-2017, A.F.), Swedish Research Council (2017-00333, A.F.), the Charles H. Revson Senior Fellowship in Biomedical Science (19-23, A.F.), National Institutes of Health (NIH) (R01MH101214, R01MH108924, R01DA050374, B.L.), the Cold Spring Harbor Laboratory and Northwell Health Affiliation (B.L.), Feil Family Neuroscience Endowment (B.L.), and German Academic Scholarship Foundation (E.C.G.).

## AUTHOR CONTRIBUTIONS

A.F. and B.L. conceived and designed the study. A.F. conducted the experiments and analyzed data. A.C. assisted with the photometry experiments with food odors and the data analysis. S. Boyle set up behavioral rigs and generated Matlab code for controlling behavioral devices and analyzing photometry data. R.S. assisted with the smFISH experiments. R.R. and J.H. assisted with operating the metabolic cages. E.C.G. assisted with the GTT and ITT experiments. R.S. and E.C.G. collected tissue samples and performed qPCR experiments. J.G. assisted with the EPM and OF experiments. S. Beyaz provided critical reagents. T.J. supervised the experiments by E.C.G. and assisted with interpreting metabolic data. S.D.S. supervised the experiments by A.C. and assisted with analyzing and interpreting the data. A.F. and B.L. wrote the paper with inputs from all authors.

## RESOURCE AVAILABILITY

### Lead Contact

Further information and requests for resources and reagents should be directed to and will be fulfilled by the Lead Contact, Bo Li.

### Data and code availability

The custom code that support the findings from this study are available from the Lead Contact upon request.

## EXPERIMENTAL MODEL AND SUBJECT DETAILS

Adult male and female mice of at least 2 months old were used for all the experiments. Mice were housed under a 12-h light/dark cycle (7 a.m. to 7 p.m. light) in groups of 2-5 animals, with food and water available *ad libitum* before being used for experiments, unless otherwise specified. All behavioral experiments were performed during the light cycle. Littermates were randomly assigned to different groups prior to experiments. All experimental procedures were approved by the Institutional Animal Care and Use Committee of Cold Spring Harbor Laboratory (CSHL) and performed in accordance with the US National Institutes of Health guidelines in an AAALACi accredited facility. The *Nts^Cre^* mouse line (Stock No: 017525) and *Ai14* (Stock No: 007908) were purchased from Jackson Laboratory. All mice were bred onto a C57BL/6J background.

## METHOD DETAILS

### Immunohistochemistry

Immunohistochemistry experiments were conducted following standard procedures (Stephenson-Jones et al., 2016). Briefly, mice were anesthetized with Euthasol (0.4 ml; Virbac, Fort Worth, Texas, USA) and transcardially perfused with 30 ml of PBS, followed by 30 ml of 4% paraformaldehyde (PFA) in PBS. Brains were extracted and further fixed in 4% PFA overnight followed by cryoprotection in a 30% PBS-buffered sucrose solution for 36-48 h at 4 °C. Coronal sections (50-μm) were cut using a freezing microtome (Leica SM 2010R, Leica). Sections were first washed in PBS (5 minutes), incubated in PBST (0.3% Triton X-100 in PBS) for 30 minutes at room temperature (RT) and then washed with PBS (3 x 5 minutes). Next, sections were blocked in 5% normal donkey serum in PBST for 30 minutes at RT and then incubated with primary antibodies overnight at 4 °C. Sections were washed with PBS (3 x 5 minutes) and incubated with fluorescent secondary antibodies at RT for 2 h. Next, sections were washed twice in PBS, incubated with DAPI (4′,6-diamidino-2-phenylindole, Invitrogen, catalogue number D1306) (0.5 µg/ml in PBS) for 5 minutes. After washing with PBS (3 x 5 minutes), sections were mounted onto slides with Fluoromount-G (eBioscience, San Diego, California, USA).

Images were taken using an LSM 710 or LSM780 confocal microscope (Carl Zeiss, Oberkochen, Germany) and visualized and processed using ImageJ and Adobe Illustrator.

The primary antibody used was chicken anti-GFP (Aves Labs, GFP1020, dilution 1:1000). Appropriate fluorophore-conjugated secondary antibodies (Life Technologies) were used to detect the primary antibodies used.

### Fluorescent *in situ* hybridization

Single molecule fluorescent *in situ* hybridization (ACDBio, RNAscope) was used to detect the expression of *Nts*, *Gad2, Slc17a6 (Vglut2)*, *cFos*, and *Cre* in the IPAC and surrounding tissues of adult mice. For tissue preparation, mice were first anesthetized under isoflurane and then decapitated. Their brain tissue was first embedded in cryomolds (Sakura Finetek, Ref 4566) filled with M-1 Embedding Matrix (Thermo Scientific, Cat. No. 1310) then quickly fresh-frozen on dry ice. The tissue was stored at -80 °C until it was sectioned with a cryostat. Cryostat-cut sections (16-μm) containing the IPAC were collected and quickly stored at -80 °C until processed. Hybridization was carried out using the RNAscope kit (ACDBio).

The day of the experiment, frozen sections were post-fixed in 4% PFA in RNAse-free PBS (hereafter referred to as PBS) at RT for 15 minutes, then washed in PBS, dehydrated using increasing concentrations of ethanol in water (50%, once; 70%, once; 100%, twice; 5 minutes each). Sections were then dried at RT and incubated with Protease IV for 30 minutes at RT. Sections were washed in PBS three times (5 minutes each) at RT, then hybridized. Probes against *Nts* (Cat. No. #420441), *Gad2* (Cat. No. #439371)*, Slc17a6 (Vglut2)* (Cat. No. #319171), *c-Fos* (Cat. No. #316921), and *Cre* (Cat. No. #312281) were applied with a 1:50 dilution to IPAC sections. Hybridization was carried out for 2 h at 40°C. After that, sections were washed twice in 1x Wash Buffer (Cat. No. 310091) (2 minutes each) at RT, then incubated with three consecutive rounds of amplification reagents (30 minutes, 15 minutes and 30 minutes, at 40°C). After each amplification step, sections were washed twice in 1x Wash Buffer (2 minutes each) at RT. Finally, fluorescence detection was carried out for 15 minutes at 40°C. Sections were then washed twice in 1x Wash Buffer, incubated with DAPI for 2 minutes, washed twice in 1x Wash Buffer (2 minutes each), then mounted onto slides with Fluoromount-G (eBioscience, San Diego, California, USA). Images were taken using an LSM 710 or LSM780 confocal microscope (Carl Zeiss, Oberkochen, Germany) and visualized and processed using ImageJ and Adobe Illustrator.

#### Detection of c-Fos with fluorescent in situ hybridization

For the mice in the food-restriction (FR) groups, food was removed at 5 p.m. the day before the testing day. Food was reintroduced to the mice 18-20 h after food-restriction (between 11 a.m. and 2 p.m.). The foods used were regular chow (PicoLab Rodent Diet 20, Cat. No. #5053*) and HFD (Bioserv HFD, Bioserv, Cat. No. # S3282). At 30 minutes after the food reintroduction, food consumption was recorded, and the mice were sacrificed. The brain tissue was processed for RNAscope. Water (Hydrogel (ClearH20)) was available ad libitum until 3 h before the mice were sacrificed. Mice in the sated group had food (regular chow) and Hydrogel (ClearH20) freely available until 3 h before the mice were sacrificed. Mice and their brain tissues in different groups underwent the experimental procedure in parallel to minimize variability.

### Viral vectors

The AAV5-Ef1a-DIO-hChR2(H134R)-eYFP and AAV9-CAG-Flex-GFP were produced by the University of North Carolina vector core facility (Chapel Hill, North Carolina, USA). The AAV9-EF1a-DIO-hChR2(H134R)-eYFP-WPRE-hGH were made by the Penn Vector Core (Philadelphia, PA, USA). The AAV2/9-CAG-DIO-TeLC-eGFP was previously described (Murray et al., 2011) and custom-packed by the Penn Vector Core (Philadelphia, PA, USA). The AAV1.Syn.Flex.GCaMP6f.WPRE.SV40, were produced by Addgene (Watertown, MA, USA). All viral vectors were aliquoted and stored at –80 °C until use.

### Stereotaxic surgery

All surgery was performed under aseptic conditions and body temperature was maintained with a heating pad. Standard surgical procedures were used for stereotaxic injection and implantation, as previously described (Stephenson-Jones et al., 2016; Zhang and Li, 2018). Briefly, mice were anesthetized with isoflurane (1–2% in a mixture with oxygen, applied at 1.0 L/minute), and head-fixed in a stereotaxic injection frame, which was linked to a digital mouse brain atlas to guide the targeting of different brain structures (Angle Two Stereotaxic System, myNeuroLab.com). Lidocaine (20 µl) was injected subcutaneously into the head and neck area as a local anesthetic.

To prepare mice for the photometry, optogenetics and inhibition experiments, we first made a small cranial window (1–2 mm^2^) in each mouse, bilaterally. We then lowered a glass micropipette (tip diameter, ∼5 μm) containing viral solution to reach the IPAC (coordinates: 0.35 mm anterior to Bregma, 1.40 mm lateral from midline, and 4.50 mm vertical from brain surface). 0.1–0.15 μL of viral solution was delivered with pressure applications (5–20 psi, 5–20 ms at 1 Hz) controlled by a Picospritzer III (General Valve) and a pulse generator (Agilent). The rate of injection was ∼20 nl/minute. The pipette was left in place for 10–15 minute following the injection, and then slowly withdrawn. Infection of the IPAC was performed in both hemispheres in mice dedicated to optogenetic and inhibition experiments, and unilaterally in mice used for photometry.

We subsequently implanted optic fibers above injection locations (coordinates: 0.35 mm anterior to Bregma, 1.40 mm lateral from midline, and 4.30 mm vertical from brain surface). A head-bar was also mounted for head-restraint. We waited for a minimum of 4 weeks following the viral injection and before starting experiments on these mice.

### *In vivo* fiber photometry and data analysis

To record the activity of IPAC^Nts^ neurons *in vivo*in behaving animals, we used a commercial fiber photometry system (Neurophotometrics Ltd., San Diego, CA, USA) to measure GCaMP6f signals in these neurons through an optical fiber (Fiber core diameter, 200 µm; Fiber length, 5.0 mm; NA, 0.37; Inper, Hangzhou, China) unilaterally implanted above the IPAC. A patch cord (fiber core diameter, 200 µm; Doric Lenses) was used to connect the photometry system with the implanted optical fiber. The intensity of the blue light (λ = 470 nm) for excitation was adjusted to ∼20 µW at the tip of the patch cord. We simultaneously recorded the isosbestic signal (using a 415 nm LED) in order to monitor potential motion artifacts as previously described (Kim et al., 2016). Emitted GCaMP6f fluorescence was bandpass filtered and focused on the sensor of a CCD camera. Mean values of fluorescent signal from each fiber were calculated and saved using Bonsai software (Bonsai), and were exported to MATLAB for further analysis. Photometry signals and relevant behavioral events were aligned based on an analogue TTL signal and timing data generated by the Bpod.

To correct photobleaching of fluorescence signals, we used a sliding window correction method to subtract the gradual reduction in baseline signal. We used Mathworks’ tsmovavg function to find the average fluorescence values calculated over a 10-second sliding time window. This gave us an average smoothed measurement of fluorescence at each timepoint throughout the bleaching process (average_window(t)) that we used for ΔF/F0 normalization, where ΔF is the change in fluorescence and F0 is baseline fluorescence. From there, we found the lowest average value in the 30 seconds before time t to get baseline fluorescence values, F0(t) = min(average_window((t– 30):t)). We treated these values as baseline fluorescence in our ΔF/F0 calculation: ΔF/F0 = (F(t) – F0(t)) / F0(t), where F is the raw fluorescence data and F0 is our normalized baseline fluorescence. This gave us a corrected ΔF/F0 that takes into account the slowly decreasing baseline and is not affected by large peaks of signal. This method of bleaching correction works very well for correcting slow bleaching and is resilient to brief disruptions in signal due to artifacts. It is not ideal for analyzing signal with large slow fluctuations, as the sliding window calculation can mask this type of signal. However, this correction method is ideal for our free-moving photometry data, since the responses to stimuli in the IPAC are transient and do not last more than the length of the 30 second sliding window. The Z-score of ΔF/F0 was then calculated using the mean and standard deviation of the signal during the baseline periods (the pooled 10 second time windows before each stimulus), Z-score(ΔF/F0) = (ΔF/F0 - mean(baseline ΔF/F0))/standard deviation(baseline ΔF/F0).

A small number of trials had artifacts due to coiling of the photometry fibers or movement of the animals. To find these trials, we searched the isosbestic control channel for large changes in fluorescence and automatically flagged for review any trial with a fluctuation (increase or decrease) of greater than 3 times the standard deviation of signals in the control channel. We discarded trials with significant artifacts during the stimulus period. This method left most trials intact, but minimized the effect of movement artifacts on the signal.

#### Photometry experiments in free moving mice (*Figure 2*)

Mice were water-restricted starting at 5 p.m. the day before the training day. On the training day, the mice learned to acquire water by licking at two adjacent spouts, with each spout delivering equal volume of water upon each lick. Water will be delivered only if mice licked a spout. The spout also served as part of a custom ‘‘lickometer’’ circuit, which registered a lick event each time a mouse completed the circuit by licking the spout. A custom software written in MATLAB (The MathWorks, Inc., Natick, Massachusetts, USA) was used to control the delivery of liquids and record licking events through a Bpod State Machine (Sanworks, Stony Brook, NY, USA) (Xiao et al., 2020). The training consisted one session of 100 trials.

The next day, which was the testing day, the mice were tested with two pairs of liquids: a sucralose solution vs. water, and a quinine solution vs. water. Each pair of liquids was available in interleaved trials (25 trials each pair, 50 trials in total; inter-trial-intervals, random between 8 and 14 s), and each liquid was delivered from one of the two spouts in equal volume (6 µl) upon each lick. The sucralose and quinine solutions were delivered from the same spout in consecutive days. The tubing and spouts were carefully washed between delivering of different liquids. Volume calibration was carried out prior to every testing.

#### Photometry experiments with olfactometer

Mice were under head-restraint in front of the output of a custom-built olfactometer. Before the testing, mice were habituated to the setup for 1 hour. The odors were presented using the olfactometer, which contains an eight-way solenoid that controls oxygen flow through eight vials. The vials contained odorants dissolved in mineral oil. The odors presented were: butyric acid (Sigma, Cat. No. #103500; 100 µl dissolved in 5 ml mineral oil), olive oil-based HFD (Envigo; 1 g in 5 ml mineral oil), coconut oil-based HFD (Envigo; 1 g in 5 ml mineral oil), white chocolate (Lindt; 1 g in 5 ml mineral oil), dark chocolate (Ghirardelli; 1 g in 5 ml mineral oil), regular chow (PicoLab Rodent Diet 20, Cat. No. #5053*; 1 g in 5 ml mineral oil), and mineral oil as a control (Sigma, Cat. No. #M3516). Food pellets were crumbled and homogenized to mineral oil for 10 minutes using a vortex mixer. Odorized oxygen was diluted 10:1 into a continuous carrier stream for a total flow of 4 l/minute. To prevent odor accumulation, air was collected behind the animal with a vacuum pump. Odor presentations were 3 s every 30 s while constantly measuring calcium signals in IPAC^Nts^ neurons. Every testing session consisted of 10 trials per odor.

For the photometry experiments with the olfactometer, we used a custom-made fiber photometry system to measure GCaMP6f signals *in vivo*. Green and red emitted fluorescence signals were filtered and split to separate photodetectors and digitally sampled at 6100 Hz via a data acquisition board (National Instruments, Model # NI USB-6211). Peaks were extracted by custom Matlab software with an effective sampling rate of 211 Hz. Signals from each signal were corrected for photobleaching by fitting the decay with a double exponential, and then normalized to a Z score. The red signals represent autofluorescence and was used to monitor and correct for potential movement artifacts (which were essentially absent as the signals were collected in head-fixed mice). The signals in the green channel were transformed back to absolute fluorescence and DF/F was computed. The resulting traces from each recording session were converted to a Z score to compare between subjects. All data analysis was performed using custom written code in Matlab.

### Liquid preference tests

To test water-restricted mice’s preference between a sucralose solution (0.13%) and water, or between a quinine solution (0.5 mM) and water (Figure 2), mice were water restricted overnight, and then were presented with two bottles, each containing one of the liquids in a pair. The mice were tested in two consecutive days. The test in each day lasted for 20 minutes, with the bottles switched their positions at 10 minutes to minimize a potential positional effect.

To test sated mice’s preference between a sucralose solution (0.004%; 0 cal/ml) and water, or between a sucrose solution (1%; 0.04 cal/ml) and water (Figure 6), sated mice were singly housed with food and water available *ad libitum* for a week before the start of the experiment. Water was dispensed through a bottle in the cage. After this, a second bottle containing either the sucralose or sucrose solution was added to the cage. Mice were allowed to first habituate to the newly added solution for 24 h, after which their consumption of the solution and water over a 48-h period was measured. The testing of sucralose and sucrose was separated by a 48-h period, during which the mice had access only to water. The positions of the bottles were switched every 24 h to minimize a potential positional effect.

### Food preference test

Mice were familiarized with HFD^CO^ and HFD^OO^ for 4 days, during which HFD^CO^ or HFD^OO^(1g/mouse) was available for 3 h in home cage on alternating days, with chow and water available *ad libitum*. The day prior to the testing, mice were singly housed. On the test day, both HFD^CO^ and HFD^OO^ were delivered to the cage and intake was measured at 3 h after the delivery.

### Optogenetics and feeding behavior

Mice sated on regular chow were habituated to the box for testing the effects of optogenetics on feeding behavior for 10 minutes on the day prior to the testing. Food was provided to the floor of the box. On the testing day, feeding behavior was assessed for 5 minutes with laser off, then 5 minutes with light on (20Hz, 7-10 mW measured at the tip of the fiber), and then another 5 minutes with laser off. Food was provided to the floor and weighed before and after each of the 5-minute sessions. The foods used were grain-based pellets (similar to the regular chow; 45 mg/pellet, Bioserv, F0165, 3.43 cal/g), sucrose (45 mg/pellet, Bioserv, F0021, 3.83 cal/g), high fat diet (HFD) (soft pellet, Bioserv, S3282, 5.49 cal/g), white chocolate (Lindt, 5.5 cal/g), dark chocolate (Ghirardelli, 5.5 cal/g), HFD^CO^ (Envigo custom diet, 4.5 cal/g), HFD^OO^ (Envigo custom diet, 4.5 cal/g). Plain and quinine-flavored grain-based pellets were prepared by immerging the pellets in either water or a 10 mM quinine solution for 10 minutes. Pellets were dried overnight and used for testing the following day. Diets were presented on consecutive days to sated mice.

To score feeding bouts, videos generated from the feeding behavioral assays were analyzed frame by frame using Behavioral Observation Research Interactive Software (BORIS) (Gamba, 2016). A feeding bout was defined as an event lasting for at least 3 seconds from pellet pickup to either pellet drop or pellet consumed.

### Real-time place aversion or preference test

Freely moving mice were initially habituated to a two-sided chamber (23 × 33 × 25 cm; made from Plexiglas) for 10 minutes, during which their baseline preference for the left or right side of the chamber was assessed. During the first test session (10 minutes), we assigned one side of the chamber (counterbalanced across mice) as the photostimulation side, and placed the mice in the non-stimulation side to start the experiment. Once the mouse entered the stimulation side, photo-stimulation (5-ms pulses, 20 Hz, 7-10 mW (measured at the tip of optic fibers)), generated by a 473-nm laser (OEM Laser Systems Inc., Bluffdale, Utah, USA), was immediately turned on. Photostimulation was turned off as soon as the mouse exited the stimulation side. In the second test session (10 minutes) we repeated this procedure but assigned the other side of the chamber as the stimulation side. The behavior of the mice was videotaped with a CCD camera interfaced with Ethovision software (Noldus Information Technologies), which was also used to control the laser stimulation and extract behavioral parameters (position, time, distance and velocity).

### Self-stimulation tests

Freely moving mice were placed in a chamber equipped with two ports. Poking into one of the ports (the active port) triggered photo-stimulation for 2 s in the IPAC (5-ms pulses, 20 Hz, 10 mW; λ = 473 nm), whereas poking into the other port (the inactive port) did not trigger photo-stimulation. Mice were allowed to freely poke the two ports and were tested in 1-h sessions.

For testing the impact of nutritional state on self-stimulation behavior, ChR2 mice were trained for two consecutive days, 1 hour per day, to nose poke on a fixed ratio (FR1) to self-stimulate IPAC^Nts^ neurons while sated on regular chow (PicoLab Rodent Diet 20, Cat. No. #5053*). Each nose poke produced a 2-second train of stimulation (5-ms pulses, 20 Hz, 10 mW; λ = 473 nm). Mice were then tested on consecutive days when fed a high fat diet (“HFD”, Bioserv, Cat. No. # S3282; Physiological value: 5.49 Kcal/g) or after being food-restricted overnight. High-fat diet was provided to mice already sated on chow for 2h prior testing. Self-stimulation experiments were carried out during the light cycle, between 9 a.m. and 5 p.m.

### Metabolic testing

Mice were singly housed and habituated to the metabolic cages (CLAMS, Columbus) for at least a week before testing, under a 12-h light/dark cycle (6 a.m. to 6 p.m. light). Mouse locomotor activity, energy expenditure (EE), oxygen consumption (VO_2_), carbon dioxide production (VCO_2_), Respiratory exchange ratio (RER), food and water intake were recorded. The mice were first fed with regular chow (PicoLab Rodent Diet 20, Cat. No. #5053*; physiological value, 3.43 kcal/g) and then with HFD (Bioserv, Cat. No. # S3282; physiological value, 5.49 kcal/g). Diets and water were available *ad libitum*. Gas sensor calibration (CO_2_, O_2_) of the apparatus was performed before each test. Mouse bodyweight was recorded prior to and after every testing session.

### Insulin tolerance test (ITT) & glucose tolerance test (GTT)

Singly housed mice were transferred to a clean cage, with food removed for 6 hours (9 a.m. – 3 p.m.) before each test. All tests started at 3 pm. For ITT, mice were injected intraperitoneally (i.p.) with 0.5 U/kg body weight insulin (Humulin, Eli Lilly; NDC Code: 0002-8215) in 0.9% sterile saline. For GTT, mice were injected i.p. with 1 g/kg bodyweight glucose (Sigma G5767-25G) in 0.9% sterile saline. There was a 48-h gap between tests, during which food and water were available *ad libitum*. Blood glucose levels were measured in duplicates at 0, 15, 30, 45, 60, 90, and 120 minutes after injection using OneTouch Ultra 2 Glucometer (OneTouch).

### RNA extraction and qPCR

Approximately 50 mg fat tissue was harvested using sterile instruments, and was frozen in 500 µl Trizol (Thermo Fisher, Cat. No. #15596026) on dry ice and stored at -80 °C until further processing. The tissue was homogenized by adding a stainless-steel bead (Qiagen, Cat. No. #69989) into each tube and shaking the tubes in the TissueLyser (TissueLyser II, Qiagen, Cat. No. #85300) 2 times for 2 minutes each at 30 Hz. After incubating the homogenate for 5 minutes on ice, 100 µl Chloroform (Sigma-Aldrich, Cat. No. #C2432-1L) was added and the tubes were shaken briefly. After incubating for 3 minutes on ice, the tubes were spun at 12000 g at 4 °C for 15 minutes. Subsequently, the clear top layer was transferred into a fresh tube and 1/10 volume of 3 M sodium acetate (Bioworld, Cat. No. #41920024-4) and Glycogen (Thermo Scientific, Cat. No. #R0551) at a final concentration of 1 µg/ul and 250 µl isopropanol (Fisher Scientific, Cat. No. #S25372) were added. The tubes were inverted to mix the contents and after 10 minutes incubation on ice, the tubes were spun at 12000 g at 4 °C for 10 minutes. The supernatant was discarded, and the RNA pellet resuspended in 500 µl 75% ethanol. After centrifuging at 7500 g at 4 °C for 5 minutes, the supernatant was discarded, and the RNA pellet left to air dry for 5 minutes and then resuspended in 25 µl RNAse-free water. cDNA was synthesized from 500 ng total RNA using Taqman Reverse Transcription reagents (Thermo Fisher, Cat. No. #N8080234). Quantitative RT-PCR was performed using QuantStudio™ 6 Flex Real-Time PCR System, using Taqman Fast Advanced Master Mix (Thermo Fisher, Cat. No. #4444556) and Taqman Primers. The 2ΔΔCt method was used to quantify relative amounts of product with a housekeeping gene (Gapdh) as endogenous control. Primers used were Gapdh (Thermo Fisher, Assay ID: Mm99999915_g1, Cat. No. #4331182) and Ucp1 (Thermo Fisher, Assay ID: Mm01244861_m1, Cat. No. #4331182).

### H&E Staining

Tissues were fixed in 4% PFA for 24 h at 4°C, washed in PBS three times at room temperature and dehydrated in 70% ethanol. Subsequently, tissues were embedded in paraffin, cut using a microtome serially to produce 5-μm sections and stained with Hematoxylin and Eosin (H&E). Pictures were taken using a Zeiss Observer microscope equipped with 10x, 20x and 40x lenses.

### Oil Red O staining

Livers were fixed in 4% PFA for 24 h at 4°C, washed in PBS three times at room temperature and cryopreserved in 30% sucrose. Tissues were embedded in OCT tissue tek (Sakura, Cat. No. #4583) and 10-μm sections were cut using a Leica Cryostat. Oil Red O staining was performed as previously described (Mehlem et al., 2013) including the counterstaining with Hematoxylin (Abcam, Cat. No. # ab220365). Pictures were taken using a Zeiss Observer microscope equipped with 10x, 20x and 40x lenses.

## QUANTIFICATION AND STATISTICAL ANALYSIS

All statistics are described where used. Statistical analyses were conducted using GraphPad Prism 7 Software (GraphPad Software, Inc., La Jolla, CA). Parametric tests were used whenever possible to test differences between two or more means. Non-parametric tests were used when data distributions were non-normal. The Shapiro-Wilk Test was used to test for normality. For two-way ANOVA tests, normality of data was assumed. All t-tests were two-tailed. Statistical hypothesis testing was conducted at a significance level of 0.05.

## FIGURES AND SUPPLEMENTARY FIGURES

**Figure S1.**
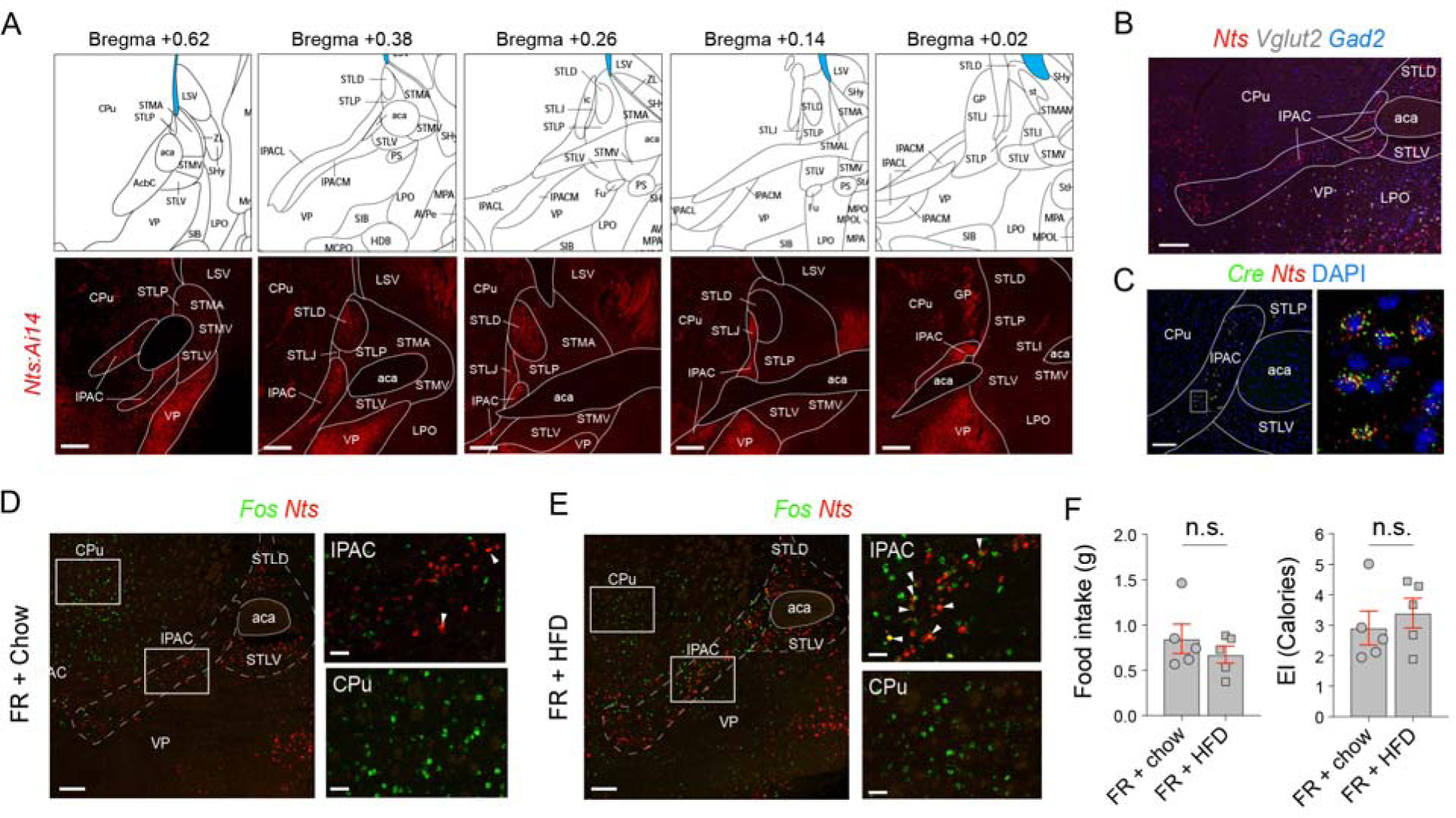
Characterization of IPAC^Nts^ Neurons, Related to Figure 1. (A) Top: coronal brain plates (from The Mouse Brain in Stereotaxic Coordinates, by Franklin and Paxinos) depicting the IPAC along the anteroposterior axis. Bottom: confocal images of coronal brain sections – which correspond to the plates on the top – from a representative *Nts^Cre^;Ai14* mouse, showing the distribution of Nts neurons in the IPAC (red). Scale bars: 200 µm. aca, anterior commissure; STLP/STLV/STLD/STLJ/STLI, lateral posterior/ventral/dorsal/juxtacapsular/intermedial nucleus of the bed nucleus of the stria terminalis; STMA/STMV, medial anterior/ventral nucleus of the bed nucleus of the stria terminalis; CPu, caudoputamen; VP, ventral pallidum; LPO, lateral preoptic area; LSV, lateral septal nucleus. (B) A representative confocal image of *in situ* hybridization for *Nts*, *Vglut2* and *Gad2* in a brain section containing the IPAC. Scale bar: 200 μm. (C) Representative confocal images of *in situ* hybridization for *Nts* and *Cre* in the brain sections containing the IPAC from *Nts^Cre^* mice. Scale bar: 100 μm. (D) Representative confocal images showing *Nts* and *cFos* expression in the IPAC and surrounding tissues in brain sections prepared from food-restricted (FR) mice just fed chow. On the right are high-magnification images of the boxed areas on the left, showing only few *Nts*^+^ cells in the IPAC expressed *cFos* (top panel, arrow heads), and many cells in the CPu expressed *cFos* (bottom panel). Scale bars: 200 μm (left panel), 50 μm (right panels). (E) Representative confocal images showing *Nts* and *cFos* expression in the IPAC and surrounding tissues in brain sections prepared from FR mice just fed HFD. On the right are high-magnification images of the boxed areas on the left, showing many *Nts*^+^ cells in the IPAC expressed *cFos* (top panel, arrow heads), and many cells in the CPu expressed *cFos* (bottom panel). Scale bars: 200 μm (left panel), 50 μm (right panels). (F) Quantification of food (left) and energy (right) intake in FR mice just fed chow or HFD (food intake, p = 0.3799 (n.s.); energy intake (EI), p = 0.5295 (n.s.); unpaired t-test).

**Figure S2.**
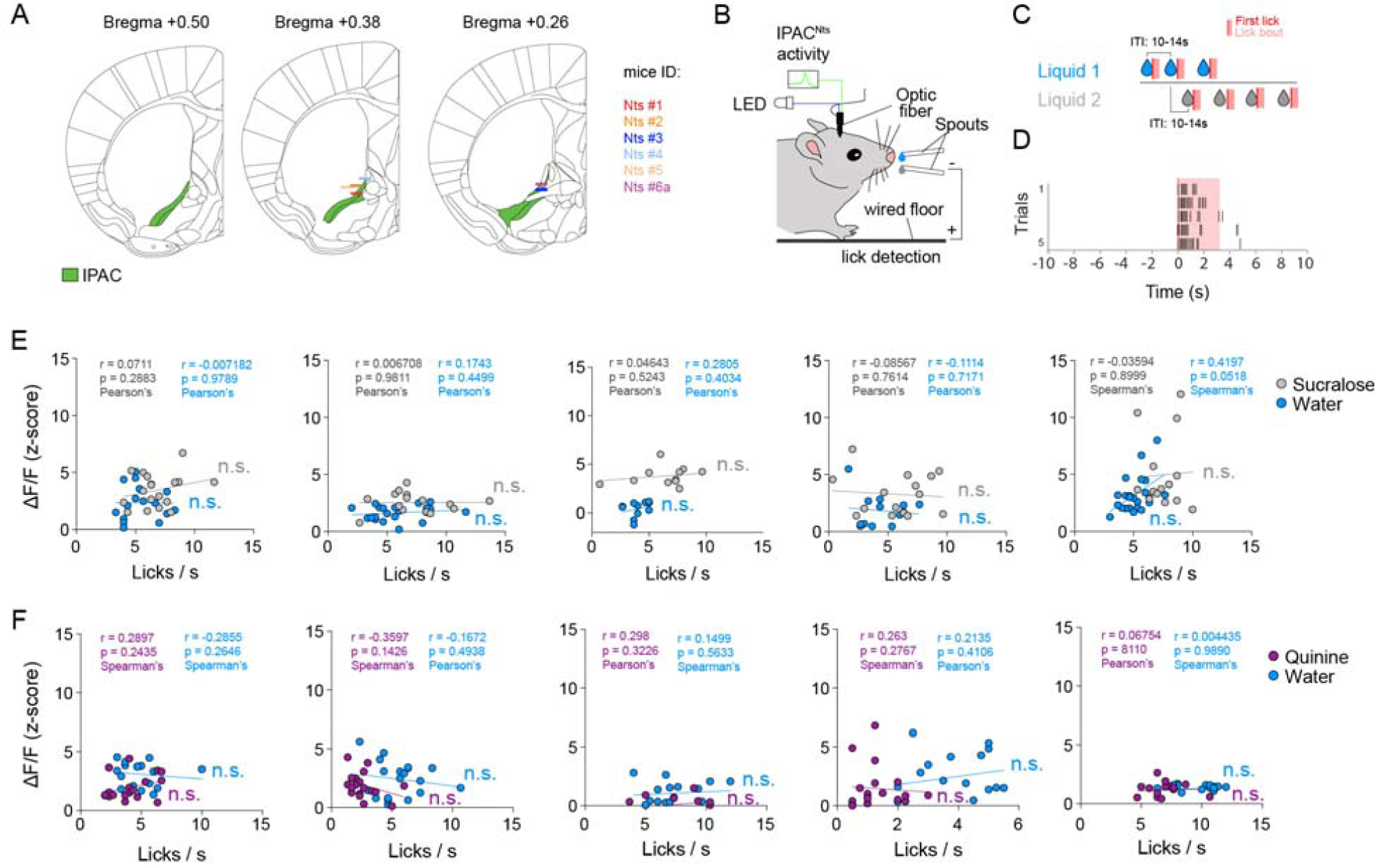
IPAC^Nts^ Neurons Do Not Represent Motion, Related to Figure 2. (A) Schematics showing the locations of optic fiber placement in the mice used in Figure 2. (B, C) Schematics of the experimental setup (B) and task structure (C). (D) Raster plot showing licking behavior following liquid delivery. (E, F) Analysis of the relationship between IPAC^Nts^ neuron responses and mouse licking behavior. Each plot represents the result from one mouse, and each dot represents data from one trial. The amplitude of peak GCaMP6 signals and average lick rate in a 3-s window following the first lick in each trial were used for the analysis. P > 0.05 (n.s.) for all mice, Pearson’s or Spearman’s correlation analysis.

**Figure S3.**
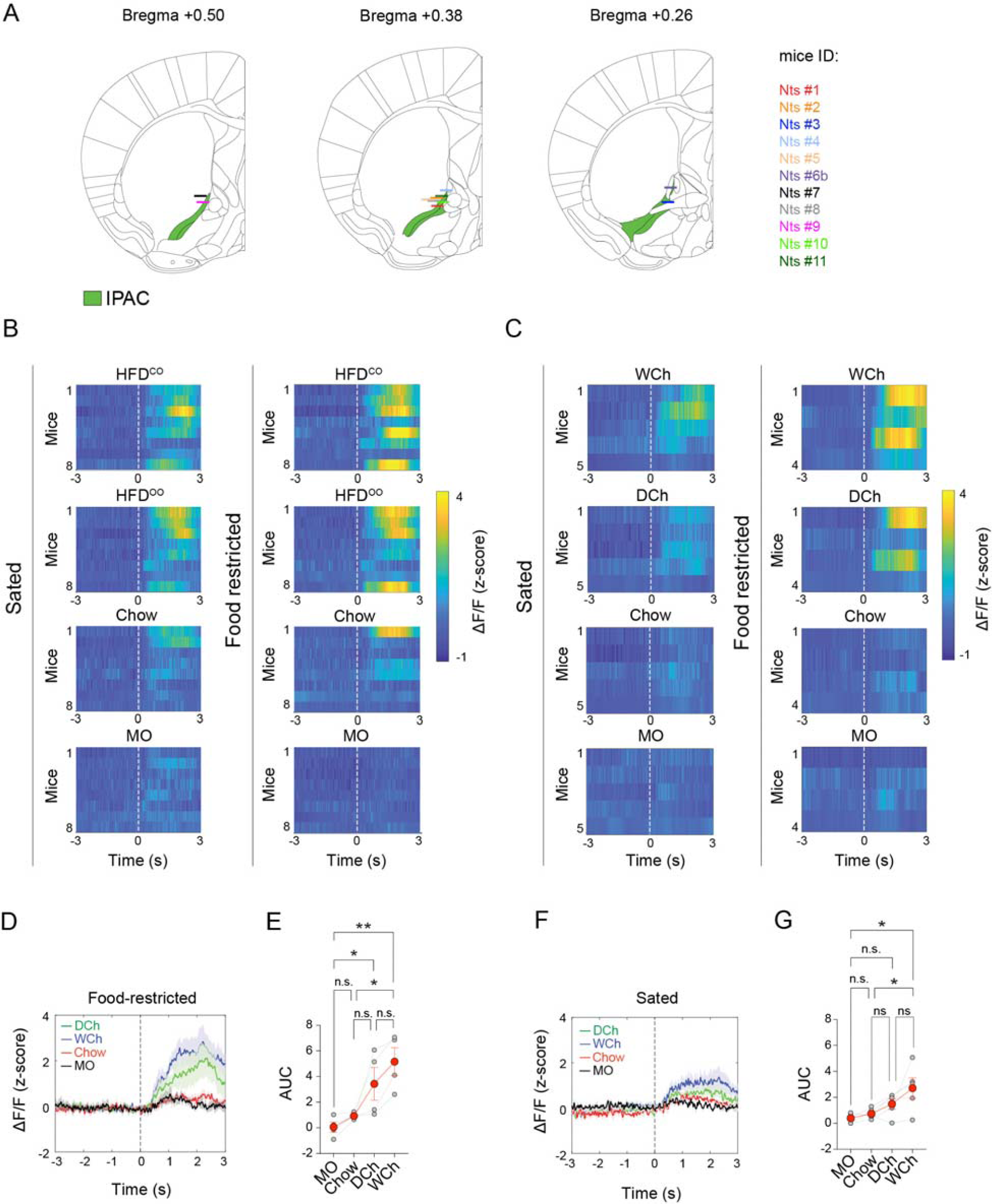
Characterization of IPAC^Nts^ Responses to Food Odors, Related to Figure 3. (A) Schematics showing the locations of optic fiber placement in the mice used in Figure 3. (B, C) Heatmaps of the response of IPAC^Nts^ neurons in individual mice to odors derived from different food sources, under sated or food-restricted condition, as indicated. Dashed lines indicate the onset of odor presentation. (D) Average GCaMP6 signals from IPAC^Nts^ neurons in food-restricted mice aligned to the presentation of different odors (dashed line). (E) Quantification of the area under the curve (AUC) of the responses in individual mice between 0 and 3 s. N = 4 mice. F_(3,9)_ = 10.36, p = 0.0028; *p < 0.05, **p < 0.01, n.s., p > 0.05; one-way RM ANOVA followed by Holm-Sidak’s multiple comparisons test. (F) Average GCaMP6 signals from IPAC^Nts^ neurons in sated mice aligned to the presentation of different odors (dashed line). (G) Quantification of the AUC of the responses in individual mice between 0 and 3 s. N = 5 mice. F_(3,12)_ = 5.169, p = 0.0160; *p < 0.05, n.s., p > 0.05; one-way RM ANOVA followed by Holm-Sidak’s multiple comparisons test.

**Figure S4.**
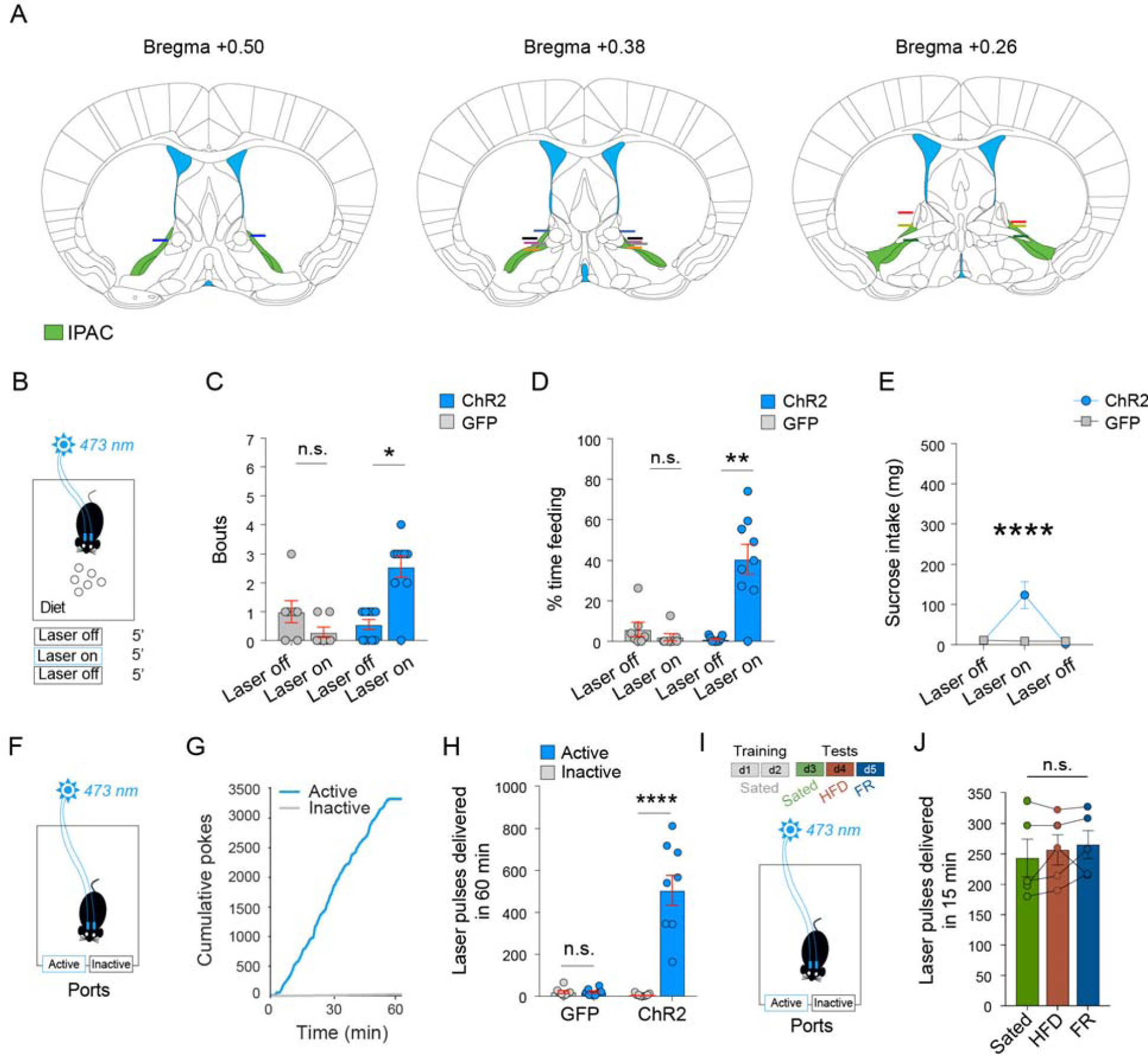
Characterization of the Effects of Activating IPAC^Nts^ Neurons, Related to Figure 4. (A) Schematics showing the locations of optic fiber placement in the mice used in Figure 4. (B) A schematic of the paradigm for testing the effects of optogenetics on feeding behavior. (C) Light delivery into the IPAC increased the number of feeding bouts in the ChR2 but not GFP mice. ChR2 mice, n = 9, *p = 0.0117; GFP mice, n = 7, p = 0.1250 (n.s.); Wilcoxon matched-pairs signed rank test. (D) Light delivery into the IPAC increased the duration of feeding bouts in the ChR2 but not GFP mice. ChR2 mice, n = 9, **p = 0.0078; GFP mice, n = 7, p = 0.1250 (n.s.); Wilcoxon matched-pairs signed rank test. (E) Light delivery increased sucrose intake in the ChR2 (n = 9), but not GFP (n = 8) mice (F_(2, 30)_ = 10.16, p = 0.0004; ****p < 0.0001; two-way RM ANOVA followed by Sidak’s multiple comparisons test). (F) A schematic of the self-stimulation paradigm. (G) Cumulative curves for the poking responses of a representative ChR2 mouse at a port where poking triggered the photostimulation (active) and a port where poking did not trigger the photostimulation (inactive). (H) Quantification of the poking responses as shown in (G). The ChR2 mice, but not the GFP mice, poked the port for photostimulation in the IPAC (ChR2 mice, n = 9, ****p = 0.0001, paired t-test; GFP mice, n = 8, p = 0.3750 (n.s.), paired Wilcoxon test). (I) A schematic of the design for testing self-stimulation under different states. (J) Quantification of the self-stimulation in the ChR2 mice under different states (n = 5 mice, F_(2,8)_ = 1.463, p = 0.2875 (n.s.), one-way ANOVA).

**Figure S5.**
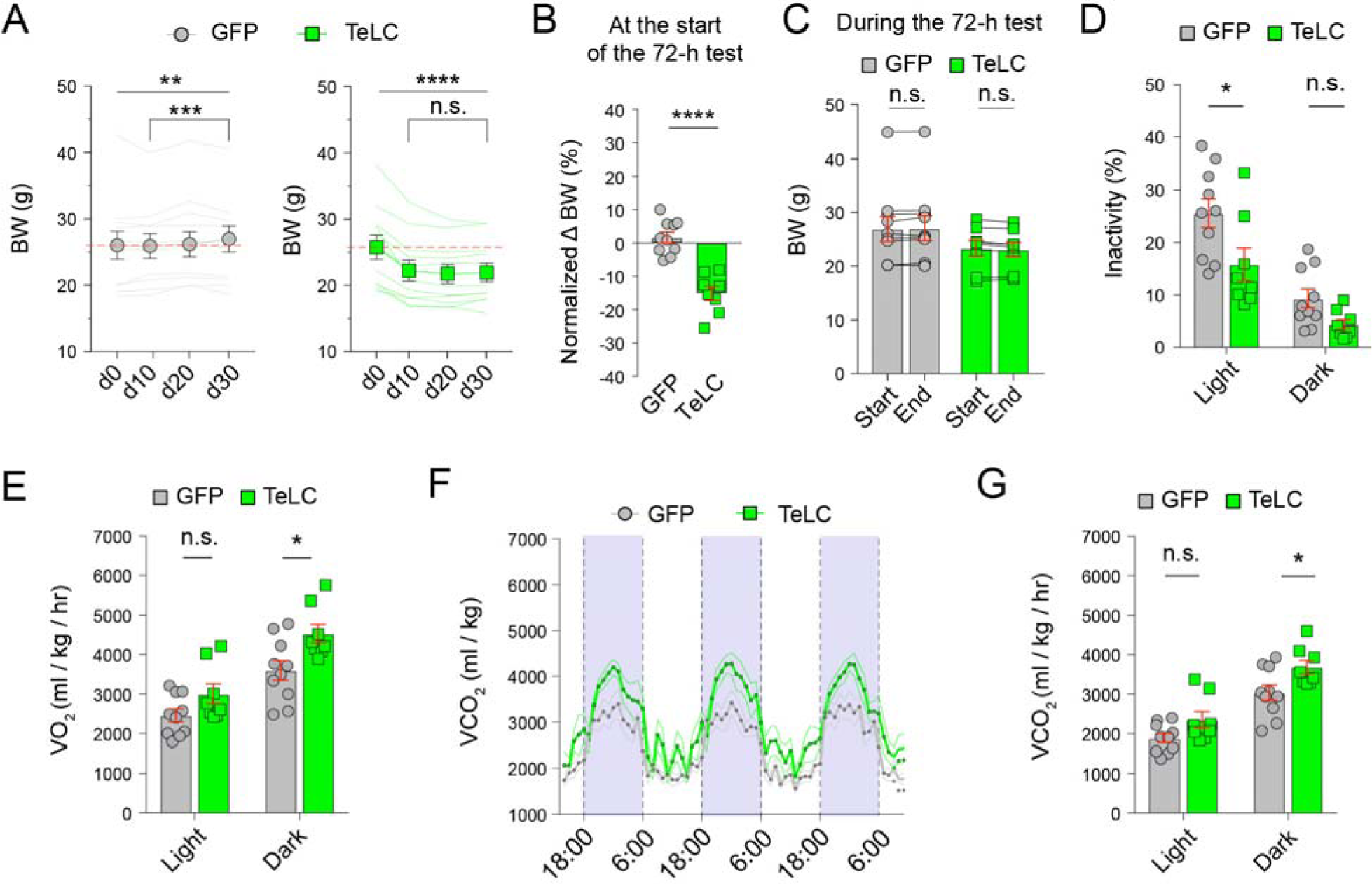
Inhibition of IPAC^Nts^ Neurons Increases Energy Expenditure, Related to Figure 5. (A) Changes in bodyweight (BW) following viral injection (d0) (GFP mice: n = 11, F_(3, 30)_ = 6.588, p = 0.0015; **p < 0.01; ***p < 0.001; TeLC mice: n = 10, F_(3, 27)_ = 28.11, p < 0.0001;****p < 0.0001, n.s., p > 0.05; one-way RM ANOVA followed by Sidak’s multiple comparisons test). (B) Changes in BW from the initial BW at d0 (GFP mice, n = 11; TeLC mice, n = 10; ****P < 0.0001, unpaired t-test). (C) BW was stable during the 72-h test (GFP mice: n = 10, p = 0.2969 (n.s.), Wilcoxon matched-pairs signed rank test; TeLC mice: n = 8, p = 0.1319 (n.s.), paired t-test). (D) Fraction of time spent without moving (inactivity) (GFP mice, n = 10; TeLC mice, n = 8; F_(1,16)_ = 6.172, p = 0.0244; *p < 0.05, n.s., p > 0.05; two-way RM ANOVA followed by Sidak’s multiple comparisons test). (E) Oxygen consumption (VO_2_) during light and dark cycles (GFP mice, n = 10; TeLC mice, n = 8; F_(1, 16)_ = 5.604, p = 0.0309; *p < 0.05, n.s., p > 0.05; two-way RM ANOVA followed by Sidak’s multiple comparisons test). (F) The volume of carbon dioxide production (VCO_2_) by GFP (n = 10) and TeLC mice (n = 8) over the 72-h period. Data are plotted in 1-h intervals. White and purple represent light (6:00-18:00) and dark cycles (18:00-6:00), respectively (F_(70, 1120)_ = 1.508, p = 0.0053, two-way RM ANOVA). (G) Carbon dioxide production (VCO_2_) during light and dark cycles (GFP mice, n = 10; TeLC mice, n = 8; F_(1, 16)_ = 5.603, p = 0.0309; *p < 0.05, n.s., p > 0.05; two-way RM ANOVA followed by Sidak’s multiple comparisons test).

**Figure S6.**
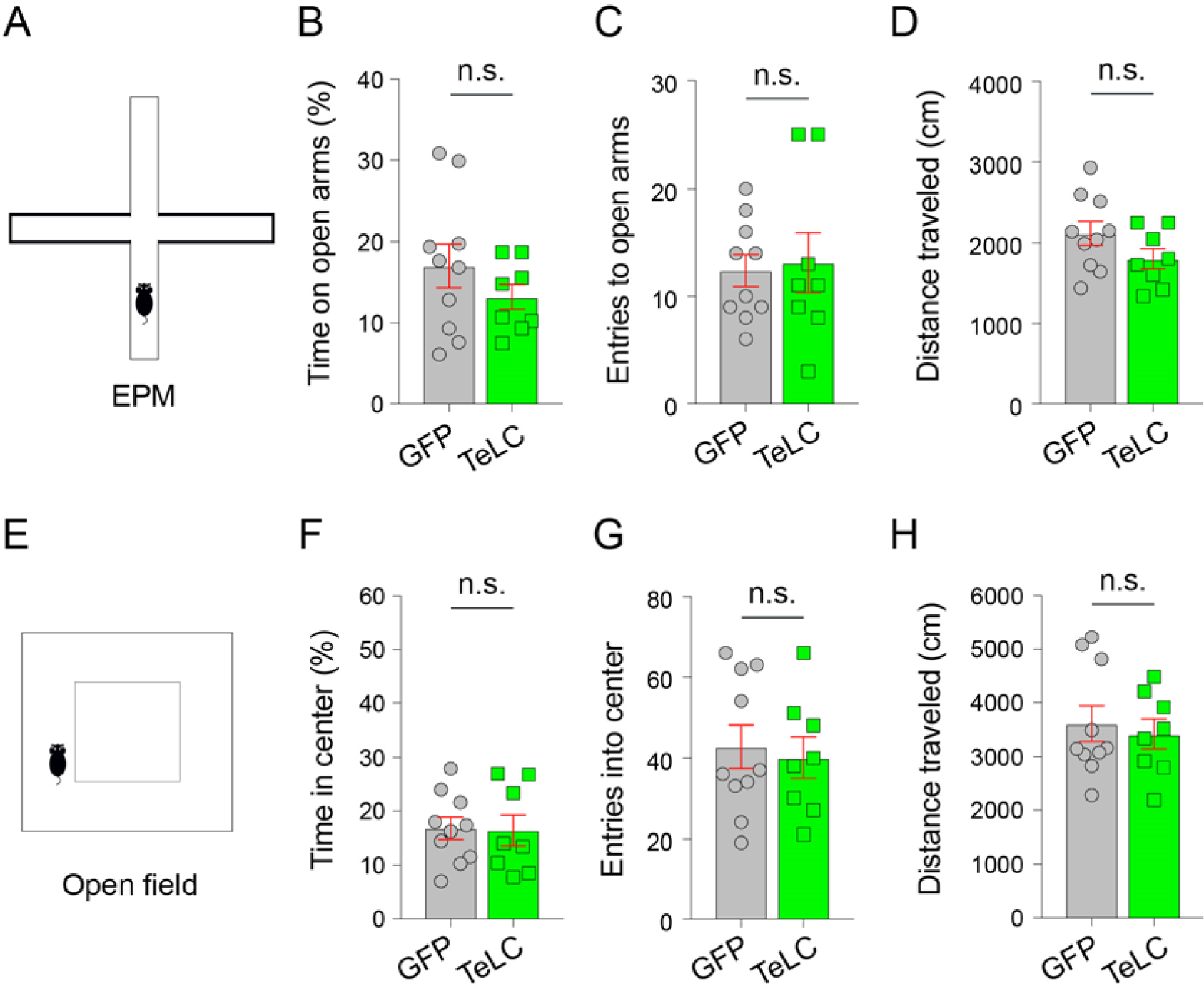
Characterization of Anxiety-Related Behaviors, Related to Figure 5. (A) A schematic of the elevated plus maze (EPM) test. (B) Time spent on the open arms (GFP mice, n = 10; TeLC mice, n = 8; n.s., p = 0.2664, unpaired t-test). (C) Entries to the open arms (n.s., p = 0.8111, unpaired t-test). (D) Distance travelled (n.s., p = 0.1345, unpaired t-test). (E) A schematic of the open field test. (F)Time spent in the center (n.s., p = 0.9068, unpaired t-test). (G) Entries into the center (n.s., p = 0.7295, unpaired t-test). (H) Distance travelled (n.s., p = 0.6640, unpaired t-test).

**Figure S7.**
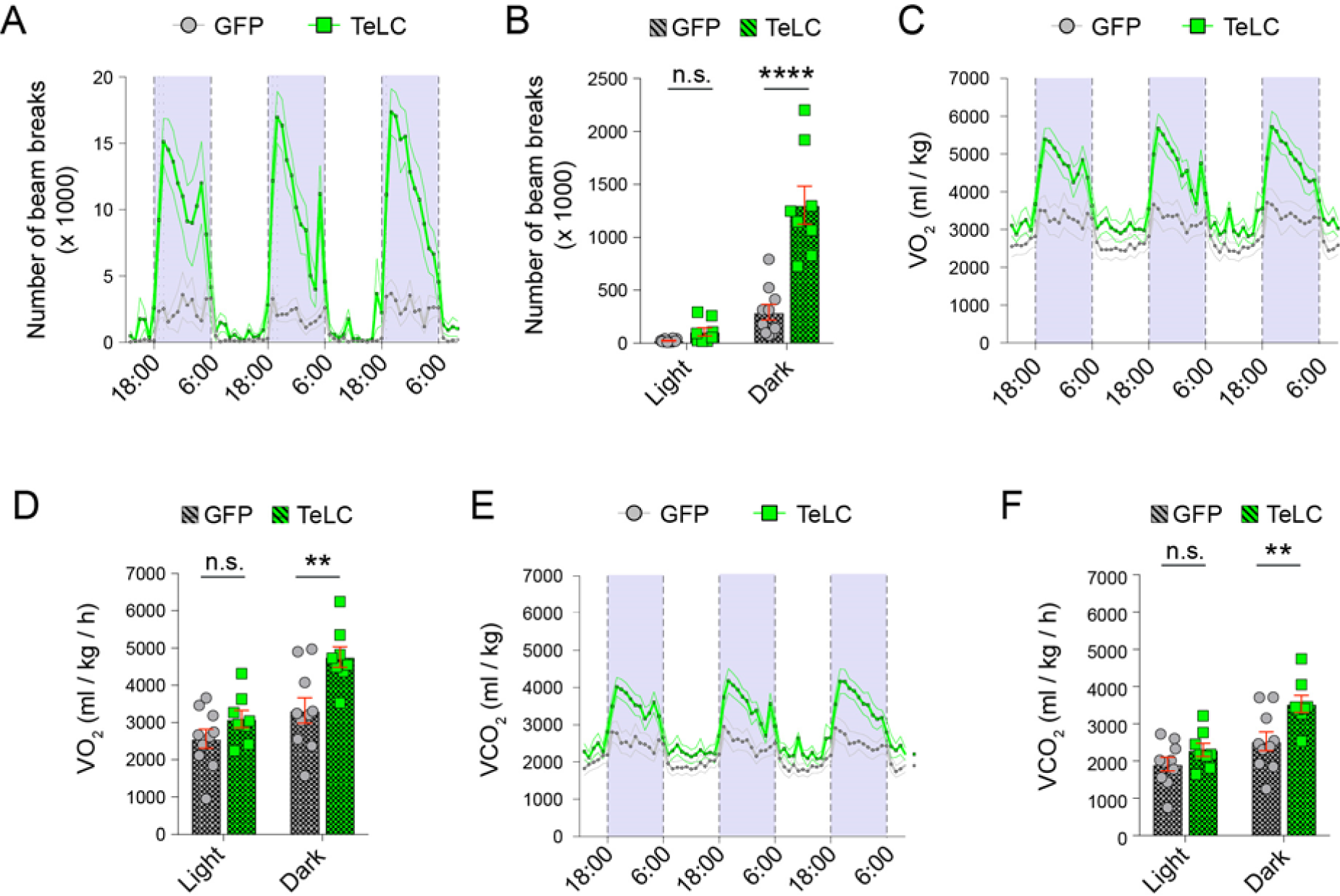
Inhibition of IPAC^Nts^ neurons in mice fed with HFD, Related to Figure 6. (A) Locomotor activity of the GFP (n = 10) and TeLC mice (n = 8) over 72 h. Data are plotted in 1-h intervals. White and purple represent light (6:00-18:00) and dark cycles (18:00-6:00), respectively (F_(71, 1136)_ = 11.46, p < 0.0001, two-way RM ANOVA). (B) Average locomotor activity of the mice in (A) during light and dark cycles (F_(1, 16)_ = 27.12, p < 0.0001; ****p < 0.0001, n.s., p > 0.05, two-way RM ANOVA followed by Sidak’s multiple comparisons test). (C) The volume of oxygen consumed (VO_2_) by GFP (n = 10) and TeLC mice (n = 8) over 72 h (F_(71, 1136)_ = 7.204, p < 0.0001, two-way RM ANOVA). (D) Oxygen consumption (VO_2_) during light and dark cycles (GFP mice, n = 10; TeLC mice, n = 8; F_(1, 16)_ = 6.05, p = 0.0257; **p < 0.01, n.s., p > 0.05; two-way RM ANOVA followed by Sidak’s multiple comparisons test). (E) The volume of carbon dioxide production (VCO_2_) by GFP (n = 10) and TeLC mice (n = 8) over the 72-h period (F_(1, 16)_ = 5.738, p < 0.0001, two-way RM ANOVA). (F) Carbon dioxide production (VCO_2_) during light and dark cycles (GFP mice, n = 10; TeLC mice, n = 8; F_(1, 16)_ = 5.276; p = 0.0355; **p < 0.01, n.s., p > 0.05; two-way RM ANOVA followed by Sidak’s multiple comparisons test).

